# Pro-inflammatory macrophages impair skeletal muscle regeneration in ischemic-damaged limbs by inducing precocious differentiation of satellite cells

**DOI:** 10.1101/2023.04.01.535211

**Authors:** Kevin W. Southerland, Yueyuan Xu, Derek T. Peters, Xiaolin Wei, Xin Lin, Yu Xiang, Kaileen Fei, Lindsey A. Olivere, Jeremy M. Morowitz, James Otto, Qunsheng Dai, Christopher D. Kontos, Yarui Diao

**Affiliations:** Division of Vascular and Endovascular Surgery, Department of Surgery, Duke University Medical Center, Durham, NC 27710, USA; Department of Cell Biology, Duke University Medical Center, Durham, NC 27710, USA; Duke Regeneration Center, Duke University Medical Center, Durham, NC 27710, USA; Center for Advanced Genomic Technologies, Duke University, Durham, NC 27708, USA; Duke University School of Medicine, Duke University, Durham, NC 27710, USA; Division of Vascular Surgery, Department of Surgery, University of Pittsburgh Medical Center, Pittsburgh, PA 15217, USA; Development and Stem Cell Biology Program, Duke University, Durham, NC 27710, USA; Division of Cardiology, Department of Medicine, Duke University Medical Center, NC 27710, USA; Department of Orthopaedic Surgery, Duke University Medical Center, Durham, NC 27710, USA; Department of Pathology, Duke University Medical Center, Durham, NC 27710, USA

**Author notes:** These authors contributed equally to this study (Kevin W. Southerland, Yueyuan Xu, Derek T. Peters).

## Abstract

Chronic limb-threatening ischemia (CLTI), representing the end-stage of peripheral arterial disease (PAD), is associated with a one-year limb amputation rate of ∼15-20% and significant mortality. A key characteristic of CLTI is the failure of the innate regenerative capacity of skeletal muscle, though the underlying mechanisms remain unclear. Here, single-cell transcriptome analysis of ischemic and non-ischemic muscle from the same CLTI patients demonstrated that ischemic-damaged tissue is enriched with pro-inflammatory macrophages. Comparable results were also observed in a murine CLTI model. Importantly, integrated analyses of both human and murine data revealed premature differentiation of muscle satellite cells (MuSCs) in damaged tissue and indications of defects in intercellular signaling communication between MuSCs and their inflammatory niche. Collectively, our research provides the first single-cell transcriptome atlases of skeletal muscle from CLTI patients and murine models, emphasizing the crucial role of macrophages and inflammation in regulating muscle regeneration in CLTI through interactions with MuSCs.

## INTRODUCTION

Atherosclerotic vascular diseases that cause tissue ischemia are the cause of pathological conditions such as myocardial infarction, stroke, and peripheral artery disease (PAD)^1–5^. In PAD, the most severe clinical manifestation is chronic limb threatening ischemia (CLTI), which is associated with a high incidence of permanent limb tissue loss ^6^. Accordingly, about 15-20% of CLTI patients undergo limb amputation within one year of diagnosis, and 50% die within five years ^7^. The current treatment options for CLTI patients focus primarily on improving limb perfusion but these strategies often fail to prevent disease progression or limb loss, pointing to a mechanism other than simply tissue perfusion as the sole etiology of tissue injury ^8^. Accumulating evidence now points to the ability of the skeletal muscle to withstand ischemic injury or to regenerate in the setting of ischemia as important mediators of tissue loss in CLTI ^9–13^. In support of this notion is the discrepancy between CLTI patients and those with intermittent claudication (IC), a mild form of PAD. (IC) presents as reproducible muscle pain with exertion that is relieved with rest ^14^. Patients with IC have more favorable clinical outcomes than CLTI patients. In fact, the one year limb loss rate for IC is <1% ^15^. Intriguingly, a subset of IC patients exhibits atherosclerosis that is comparable to those observed in CLTI but does not develop the permanent tissue loss phenotype characteristic of CLTI (**Fig S1A**) ^14,16^. The distinct clinical outcomes of patients with IC versus CLTI suggest that the reparative capacity of skeletal muscle plays a key role in determining the difference in disease progression in the two groups. Indeed, CLTI limbs exhibit extensive fibrosis and fatty deposition in skeletal muscle and a distinct skeletal muscle mitochondriopathy that distinguishes it from IC, indicating a role for pathologic alteration in skeletal muscle regeneration in the development of CLTI ^17,18^. Therefore, a thorough understanding of the cellular and molecular mechanisms underlying ischemia-induced muscle regeneration will likely shed light on the development of new regenerative medicine strategies for limb salvage in CLTI patients that are independent of limb perfusion.

In a pre-clinical murine model of CLTI, in which ligation of the femoral artery causes hind limb ischemia (HLI), the degree of tissue loss is highly strain-dependent ^19^. Specifically, BALB/c mice develop an extensive and irreversible limb tissue loss phenotype while C57BL/6 (BL6) mice are resistant to tissue loss and can initiate a potent muscle regeneration program ^13^. Thus, murine models of HLI provide a unique opportunity to study the mechanistic determinants of skeletal muscle regeneration in the context of ischemia. In mammals, successful skeletal muscle regeneration requires the orchestrated activation, proliferation, and differentiation of muscle satellite cells (MuSCs, also known as muscle stem cells) that are normally quiescent in the uninjured state ^20,21^. The regenerative capacity of MuSCs is supported by various cell types that comprise the MuSC niche, including macrophages ^22–25^. Upon muscle injury, pro-inflammatory macrophages are enriched at the site of tissue damage, followed by a polarization process in which they acquire an anti-inflammatory and pro-reparative state in later stages of tissue repair ^26,27^. Inflammatory macrophages and their subsequent polarization to a regenerative phenotype play a critical role in numerous aspects of muscle repair and regeneration, including extracellular matrix (ECM) assembly, phagocytosis of tissue debris, and angiogenesis, in addition to supporting the proliferation and differentiation of MuSCs through the secretion of pro-regenerative factors ^22^. Despite the well-established role of macrophages in skeletal muscle regeneration, whether and how macrophages regulate limb tissue repair in response to an ischemic insult in the context of CLTI remains to be elucidated.

Here, through single-cell transcriptomic analysis of skeletal muscle tissue from representative CLTI patients and murine models of HLI, we show that non-regenerative, ischemia-injured limbs are enriched with macrophages exhibiting a persistent pro-inflammatory signature. Significantly, these pro-inflammatory macrophages do not express the pro-regenerative cytokines that normally promote the proper balance in MuSCs of proliferation versus myogenic differentiation. Our findings support the idea that macrophages play a critical role in regulating limb regeneration in ischemia-induced tissue damage and they provide the first single-cell transcriptome atlas of CLTI, both in humans and mice, as a valuable resource for future studies of CLTI pathobiology and potential regenerative therapeutics.

## RESULTS

### The ischemia-injured muscle in CLTI patients is enriched with pro-inflammatory macrophages

To understand the pathological changes in ischemia-injured skeletal muscle, we carried out single-cell RNA-seq analysis of fresh skeletal muscle samples from CLTI patients undergoing limb amputation surgery (**Fig 1A**). The biopsies were taken from both the distal (ischemic) and proximal (non-ischemic) regions of the amputated limb, disassociated into single cell suspensions and subjected to single-cell transcriptome analysis (**Fig 1A, Fig S1A**). Importantly, obtaining matched proximal and distal tissue from the same individual allowed us to examine the specific pathological changes caused by ischemia while controlling for differences in the genetic background and health conditions of each patient. Using pairs of matched proximal and distal muscle tissues from three representative CLTI patients, we recovered a total of 16,201 high-quality cells for downstream bioinformatics analysis. After correcting the batch effect and patient-specific biases (detailed in Materials and Methods), all the cells were distributed across the major cell clusters on the UMAP plot (**Fig S1B**). They were annotated into eight major cell types (**Fig 1B**), including fibro-adipogenic progenitor cells (FAPs), MuSCs, myoblasts, endothelial cells, macrophages, and neutrophils, based on the expression of well-defined cell type-specific marker genes (**Fig 1C, Fig S1C**).

**Figure 1.**
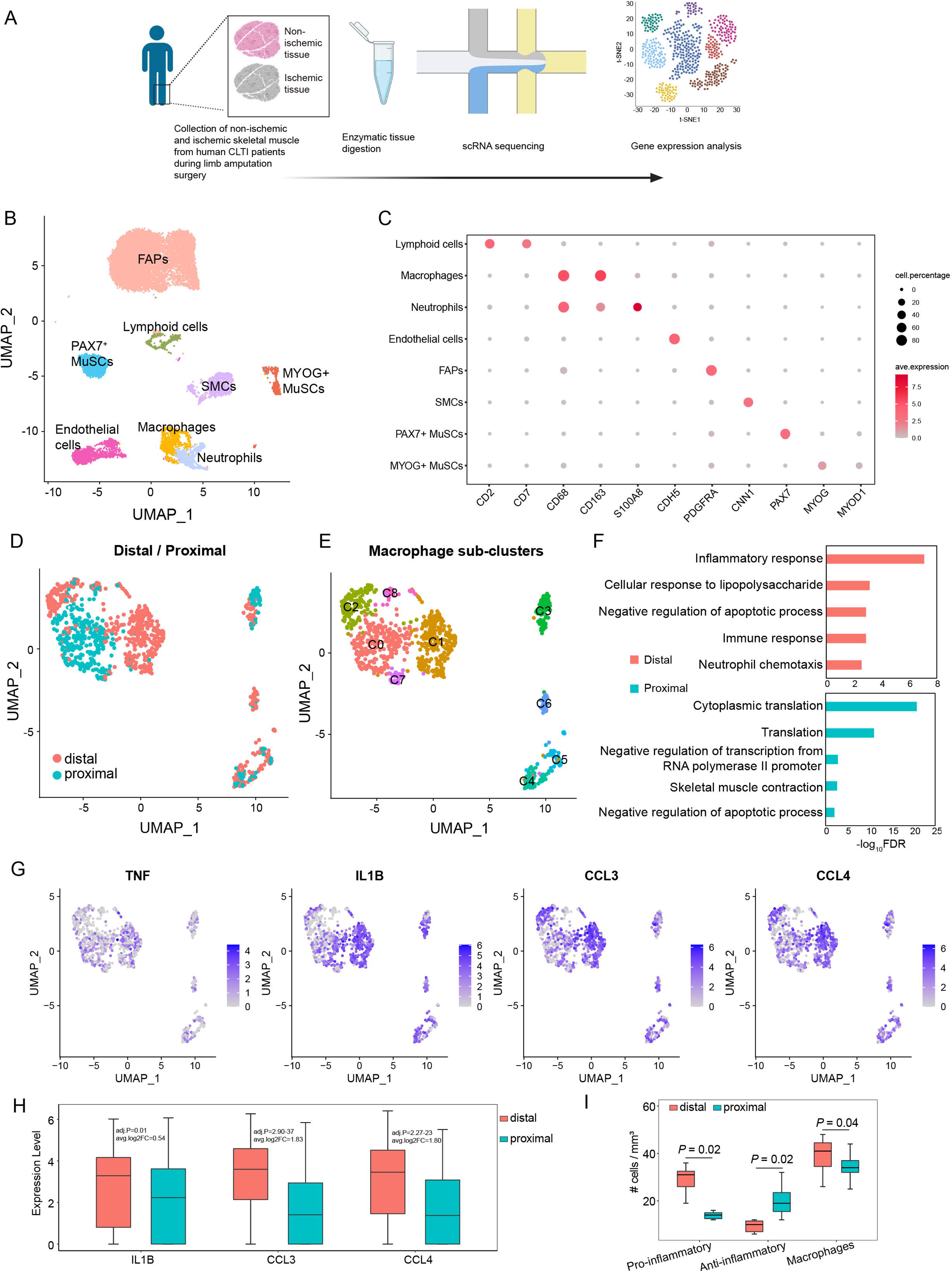
Single-cell transcriptional profiling of human CLTI patients’ limb muscle in non-ischemic versus ischemic conditions. **A:** Schematic diagram illustrating the generation of scRNA-seq datasets using proximal and distal tissue from human CLTI skeletal muscle. **B:** Uniform Manifold Approximation Projection (UMAP) visualization showing cell populations (n=16,201) from non-ischemic and ischemic tissues of CLTI patients (n=3 donors, paired proximal and distal tissues were analyzed). **C:** Dot plot displaying the expression of marker genes for each cell population. Dot size represents the percentage of cells that positively detect the transcripts, and the color scale indicates average expression levels. **D-E:** UMAP visualization of macrophages in non-ischemic and ischemic skeletal muscle, colored by condition (D, red for distal, blue for proximal) and sub-clusters (E, C0-C8). **F:** Top five GO terms enriched by differentially expressed genes (P-value < 0.05 & log2FoldChange > 0.25) between ischemic-specific clusters (1 and 2) and non-ischemic-specific cluster (0). Red and blue bars represent the GO terms enriched in distal and proximal conditions, respectively. G: Feature plots showing the expression of pro-inflammatory genes in macrophages. **H:** Quantification of representative pro-inflammatory gene expression in proximal versus distal macrophages. **I:** Quantification of CD11b+/CD206+ and CD11b+/CD206-macrophages in ischemic and non-ischemic CLTI patient muscle specimens. Data are expressed as mean ± SEM. *p ≤ 0.05.

Intriguingly, when macrophages were segregated into non-overlapping populations on a new UMAP space with increased resolution, significant differences were seen between cells derived from proximal versus distal tissue (**Fig 1D**). The macrophages were separated into nine sub-clusters (**Fig 1E**). Of these, cluster 0 was comprised primarily of cells from non-ischemic-tissue, while clusters 1 and 2 were predominantly composed of macrophages from ischemia-injured distal muscle (**Fig 1E**). We further identified genes differentially expressed in clusters 1 and 2 versus cluster 0 (Wilcoxon test, p-value < 0.05, log2 fold change > 0.25, **Table S1**) and found that the genes highly expressed in clusters 1 and 2 were enriched for Gene Ontology (GO) terms related to pro-inflammatory pathways (**Fig 1F**). Indeed, several well-characterized pro-inflammatory genes, such as TNF, IL1B, CCL3, and CCL4 ^28^, were expressed at significantly higher levels in macrophages from ischemic versus those from non-ischemic muscle (**Fig 1G, 1H**). These results demonstrate that macrophages in the ischemia-injured limb muscle of CLTI patients display a pro-inflammatory phenotype.

To validate these findings, we collected distal and proximal skeletal muscle samples from another seven CLTI patients and immunostained tissue sections with antibodies against the pan-macrophage marker CD11b and the anti-inflammatory macrophage marker CD206 (**Fig S1D**). According to previously published criteria ^29,30^, we designated the CD11b+/CD206+ cells as anti-inflammatory macrophages, and the CD11b+/CD206-cells as inflammatory macrophages. In the ischemia-injured distal muscle, we found average 1.97-fold more pro-inflammatory macrophages compared to the non-ischemic condition (**Fig 1I,** P value = 0.02, paired samples Wilcoxon test). In contrast, there were average 2.07-fold more anti-inflammatory macrophages in the non-ischemic proximal muscles (**Fig 1I,** P value = 0.02, paired samples Wilcoxon test). Considering all these results, we conclude that pro-inflammatory macrophages are enriched in the ischemia-damaged limb muscle of CLTI patients.

### Single-cell transcriptomic analysis of regenerative versus CLTI-like mouse limb muscle following hind-limb ischemia surgery

Next, we sought to determine whether the temporal change of the inflammatory response of ischemic limb muscle is associated with the tissue loss phenotype observed in the CLTI patients. Since it is not feasible to obtain clinical samples from CLTI patients over time, we employed a murine model of CLTI in which hind-limb ischemia (HLI) surgery is used to ligate the femoral artery in BALB/c and C57BL/6 mice to assess the temporal dynamics of tissue loss in CLTI (**Fig 2A, 2B**). As noted, following HLI, BALB/c mice develop a CLTI-like, profound tissue loss phenotype and paw necrosis (**Fig 2B**) ^13,19,31^. In contrast, C57BL/6 mice display very minor, if any, tissue loss (**Fig 2A, 2B**), despite experiencing a similar 80-90% reduction in limb blood flow after HLI (**Fig 2B,** left). Consistent with previous results ^13^, the ischemic tibialis anterior (TA) muscle of C57BL/6 mice exhibited a potent regenerative response following HLI, indicated by 6.2-fold greater expression of embryonic myosin heavy chain (eMHC) and 3.1-fold more Pax7+ satellite cells compared to BALB/c mice at 7 days post-injury (dpi) (**Fig 2C, 2D**). Therefore, BALB/c mice represent a murine model of CLTI with permanent tissue loss, whereas C57BL/6 mice are resistant to ischemia-induced muscle loss and display a potent skeletal muscle regenerative program following HLI.

**Figure 2.**
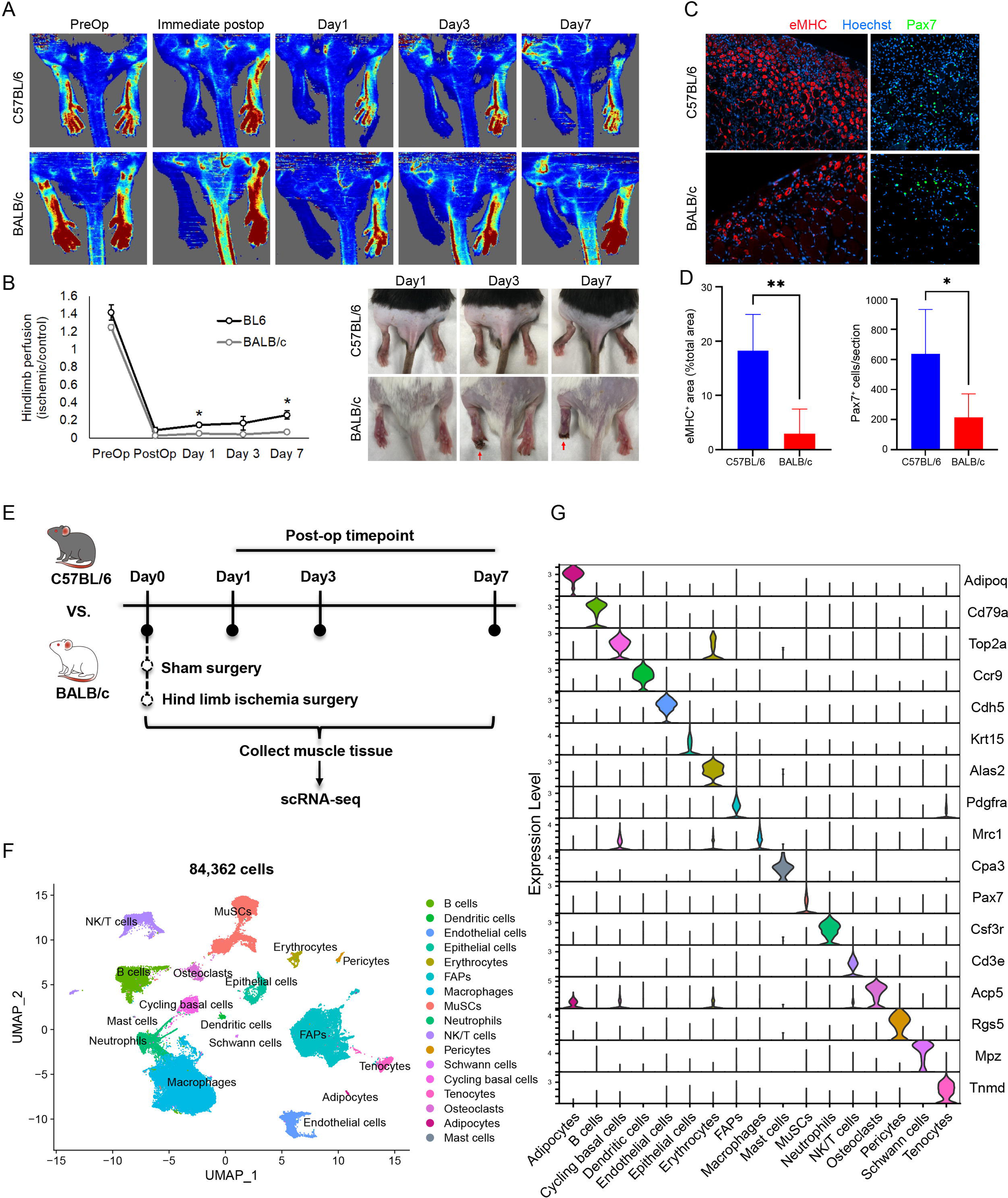
Single-cell RNA-seq analysis of hind limb ischemia (HLI) surgery induced limb muscle regeneration and damage responses in C57BL/6 and BALB/c mice. **A:** Perfusion imaging of C57BL/6 (top) and BALB/c (bottom) mouse strains before and after HLI surgery (Pre-operatively and Post-operatively, respectively) and on post-op days 1, 3, and 7. **B:** (Left) Quantification of limb perfusion as determined by perfusion imaging at indicated timepoints. Left hindlimb (HLI surgery) perfusion normalized to right hindlimb perfusion for each mouse (n=3 per mouse strain, per timepoint). *p<0.01. (Right) Representative images of mice on post-op days 1, 3, and 7 following HLI surgery. Red arrow indicates ischemic changes apparent on post-op days 3 and 7 in BALB/c mice. **C:** Representative immunofluorescence staining images of mice on post-op day 7 following HLI surgery. eMHC (red) indicates regenerated muscle fibers; Pax7 (green) indicates satellite cells. **D**: Quantification of eMHC+ area (Left panel) and Pax7+ cell counts (Right panel) shown in (C). Data expressed as mean ± SEM. *p≤0.05, **p≤0.01. **E**: Experimental design of mice scRNA-seq analysis of mouse models of HLI (n=2 per mouse strain, per timepoint). **F:** UMAP visualization of the scRNA-seq atlas assembled from all samples and time points. **G:** The expression of cell type marker genes used for each cell type/cluster annotated in (F).

To determine the temporal dynamics of inflammatory response upon limb ischemia injury, we conducted scRNA-seq analysis using cells prepared from three hindlimb muscles (TA, gastrocnemius, and the soleus) from both BALB/c and C57BL/6 mice before and after HLI at intervals from 1-7 dpi. We included two biological replicates for each experimental condition (**Fig 2E**). In total, we recovered 84,362 high-quality single cells from the two mouse strains at four-times (**Fig 2F, S2A, S2B**). We identified 17 major cell types, including muscle progenitor cells, immune cells, and FAPs (**Fig 2F**). These annotated cell types express high levels of expected marker genes that were defined in previous studies of scRNA-seq analysis of mouse skeletal muscle regeneration (**Fig 2G, S2C**). Notably, these scRNA-seq datasets thus provide the first reference atlas to examine the temporal dynamics of cell populations and gene expression patterns in mouse strains that display either successful or failed skeletal muscle regeneration following limb ischemia.

### Pro-inflammatory macrophages are enriched in the ischemic-damaged limb muscle of mice subjected to HLI

We next explored whether differences exist in the macrophage populations in the ischemic hind limbs of C57BL/6 and BALB/c mice. Fine resolution sub-clustering analysis of a total of 26,991 macrophages revealed 12 sub-clusters (**Fig S3A**, clusters 0-11), which display temporal-, strain-specific-, and cluster specific gene expression patterns (**Fig 3A, Fig S3B, Table S2**). Notably, clusters 4 and 5 were dominated by BALB/c cells, while clusters 1, 2, 3, 7, 8, and 9 were made up primarily of C57BL/6 macrophages (**Fig 3A,** right). At 3 dpi, the macrophages from BALB/c and C57BL/6 were segregated into two non-overlapping populations on the UMAP (**Fig 3A,** right, day 3), indicating drastically different gene expression programs of macrophages at day 3 between the two strains. Significantly, the BALB/c specific cluster 5 displayed a strong pro-inflammatory gene expression signature, indicated by a high “inflammatory response score” computed by the expression level of a list of genes included in the GO term of the inflammatory response (**Fig 3B**, detailed in material and methods). Macrophages can be classified largely into the pro-inflammatory M1 and anti-inflammatory/pro-regenerative M2 states based on their in vitro signatures in response to inflammation ^32^. Using this convention, we found that the anti-inflammatory M2 gene sets were highly expressed in the C57BL/6 macrophages, while the pro-inflammatory M1 gene sets were highly expressed in macrophages from BALB/c mice (**Fig 3C**). Indeed, the well-characterized anti- and pro-inflammatory genes were highly expressed in the 3dpi macrophages of C57BL/6 and BALB/c, respectively (**Fig 3D**). These results suggest that following HLI, the non-regenerative BALB/c limb muscles are enriched with macrophages exhibiting a pro-inflammatory phenotype compared to those in C57BL/6 muscle.

**Figure 3.**
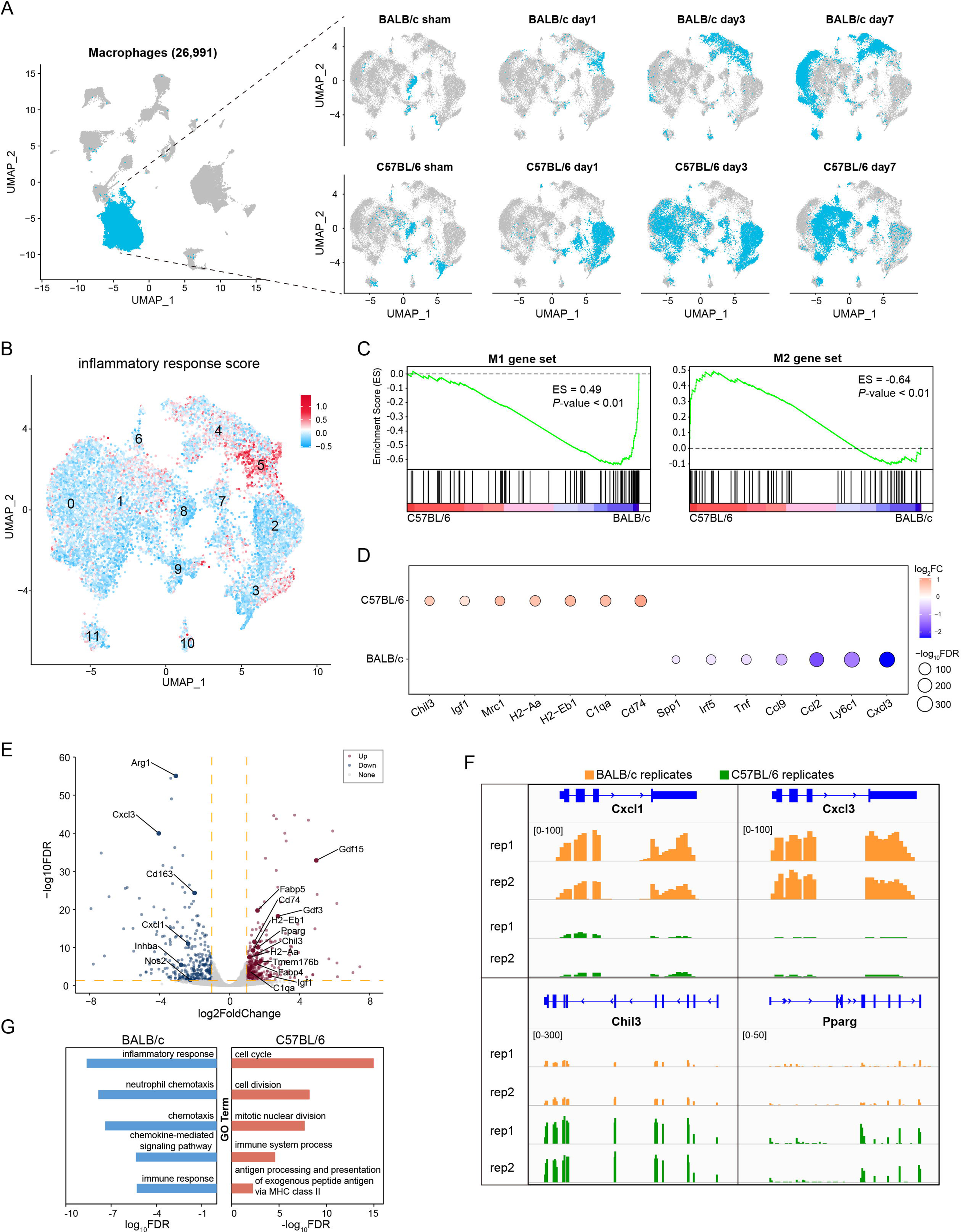
scRNA-seq analyses reveals that the pro-inflammatory macrophages are enriched in the BALB/c limb following HLI surgery. A: UMAP visualization of all cells (left) and macrophages (right) from C57BL/6 and BALB/c mice at indicated time. (left) The UMAP contains all the mouse cells from all time points in two strains. Blue cells: macrophages. B: The inflammatory gene module score is high in cluster 5 cells, which are specific to BALB/c mice. C: GSEA enrichment analysis reveals that M1 macrophage markers are highly expressed in BALB/c macrophages, while M2 macrophage markers are highly expressed in C57BL/6 macrophages. Macrophage gene expression patterns are assessed using scRNA-seq data. D: Dot plot showing differentially expressed genes in macrophages between C57BL/6 and BALB/c mice on day 3 post-ischemia. Dot size represents −log10 FDR; color scale indicates log2-fold change in gene expression. E: Volcano plot displaying differentially expressed genes from bulk RNA-seq analysis of macrophages purified from BALB/c and C57BL/6 mice at 3 days post-HLI. Red and blue dots represent upregulated genes in C57BL/6 and BALB/c mice, respectively. F: Genome browser views of bulk RNA-seq data at the indicated gene loci in macrophages from both strains on day 3. Numbers in brackets indicate the range of signal intensities. G: Top five GO terms enriched by differentially expressed genes (FDR<0.05, log2FoldChange>1) from panel E. Red and blue bars correspond to C57BL/6 and BALB/c mice, respectively.

To further validate the findings from scRNA-seq, we purified CD11b+/F480+ macrophages by FACS from the hindlimb muscle of C57BL/6 and BALB/c mice at 3-days post HLI for bulk RNA-seq analysis (**Fig S3C**). We identified 289 down-regulated and 320 up-regulated genes in C57BL/6 versus BALB/c macrophages (**Fig 3E, Table S3,** DEseq2, fold change 2, FDR 0.05). Many pro-inflammatory genes, such as *Arg1, Cxcl3*, and *Cxcl1* ^27^, were highly expressed in the BALB/c macrophages (**Fig 3E, 3F**). In contrast, anti-inflammatory and pro-regenerative genes, such as *Chil3, Igf1*, Gdf15, and *Gdf3* ^33,34^, were highly expressed in C57BL/6 3dpi macrophages (**Fig 3E, 3F, S3D**). The genes highly expressed in 3dpi BALB/c macrophages were enriched for GO terms associated with inflammatory response, chemotaxis, and immune response (**Fig 3G**). Collectively, these results highlight the clear transcriptional differences in macrophages present in the regenerative (C57BL/6) and non-regenerative (BALB/c) limbs following HLI. Furthermore, these findings demonstrate that BALB/c macrophages display a strong pro-inflammatory transcriptional signature associated with permanent limb tissue loss phenotype.

### MuSCs in the ischemia-injured limbs of BALB/c mice undergo precocious myogenic differentiation

MuSCs directly contribute to muscle regeneration thought their activation, proliferation, differentiation, and fusion processes from the initial quiescent state ^35^. To delineate strain-specific differences in MuSCs responses following HLI, we performed an in-depth analysis of all the scRNA-seq data on *Pax7*+ MuSCs and *Myod*+ MuSC-derived muscle precursor cells (MPCs). First, the MuSCs/MPCs were classified into quiescent (*Pax7* high), activated/proliferating (*MyoD, Ki67* high), and early (*MyoG* +) and late (*Ckm*+) differentiating states (**Fig 4A, S4A**). Next, we conducted pseudotime trajectory analysis to rank all the MuSCs/MPCs based on their transcriptome similarities (**Fig S4B**). This approach showed that the MuSCs/MPCs ranked at early pseudotimes expressed a high level of quiescence marker genes (*Hes1, Calcr, Cd34, Pax7, Myf5*, and *Notch1/3*), while the cells at later times, defined as those in activation/proliferation (covered by blue bar on the left) and differentiation (covered by red bar on the left) states, expressed high levels of the activation marker *MyoD*, cell cycle-related genes (*Cdnb1/2, Cdc20, Cdk1*), and myogenic differentiation markers (*MyoG, Ckm*, and *Myh1*) (**Fig 4B**). Significantly, along the pseudotime trajectory, we noted that MuSCs from both BALB/c and C57B/6 mice can undergo activation/proliferation (**Fig 4C**, blue) and differentiation (**Fig 4C**, pink) from the initial quiescent state (**Fig 4C**, yellow), demonstrating that BALB/c mice do not lack intrinsic MuSC regenerative capacity.

**Figure 4.**
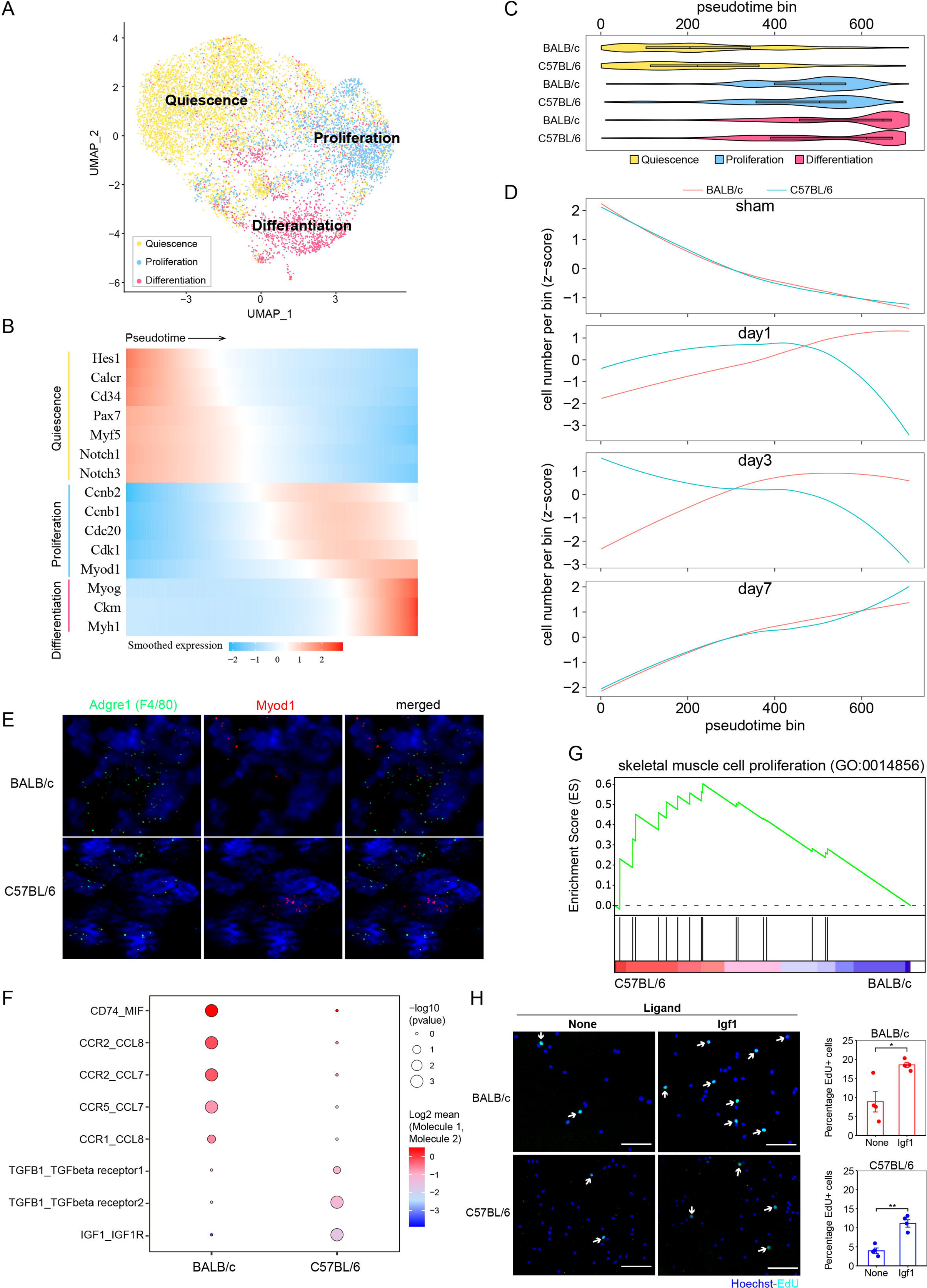
MuSCs/MPCs in BALB/c mice undergo precocious differentiation after HLI surgery. **A:** UMAP visualization of muscle stem cells (MuSCs) and muscle precursor cells (MPC) that are quiescent (yellow), activated/proliferative (blue), and differentiating (pink). Total MuSC/MPC, n=9217. **B:** All MuSCs/MPCs shown in (A) are ranked along a pseudotime trajectory, starting from quiescent cells progressing towards activated/proliferative and differentiating cells. The color of the heatmap indicates the expression level of the indicated genes in MuSC/MPCs aligned along the pseudotime. **C:** Violin plot showing the distribution of quiescent (yellow), activated/proliferative (blue), and differentiating (pink) MuSCs/MPCs in the two strains along the pseudotime trajectory shown in (B). **D:** Curve plot showing the distribution of MuSC/MPCs at each time point (before and after HLI surgery) in the two mouse strains along the pseudotime trajectory shown in (B). **E:** RNAscope data showing that Adgre1+ macrophages (F4/80, green) and Myod1+ MuSC/MPCs (red) are spatially proximal to each other in the limb muscle of BALB/c and C57BL/6 mice at day 3 after HLI. **F:** Inferred ligand-receptor interactions between macrophages and MuSCs in BALB/c and C57BL/6 following HLI at 3 days post HLI surgery. **G:** GSEA plot of gene set related to skeletal muscle cell proliferation between C57BL/6 and BALB/c mice. **H:** IGF1 promotes proliferation of primary MPCs purified from BALB/c and C57BL/6 strains. (Left) Representative images of EdU incorporation by C57BL/6 and BALB/c primary MPCs cultured with or without recombinant IGF1 for 72 hours. EdU was added to the culture medium 6 hours prior to cell fixation and imaging. Nuclei stained with Hoechst. Arrows indicate EdU+ cells. (Right) Quantification of the percentage of EdU+ cells for the indicated strains and treatment conditions. *P<0.05, **P<0.005.

It is well established that the proliferation phase of MuSCs/MPCs is critical for efficient muscle regeneration by producing sufficient MPCs for myogenic differentiation and fusion, and that the timing of the transition between proliferation and differentiation is important ^36^. We found that before HLI (day 0) and at 7dpi, the MuSCs/MPCs collected from the two mouse strains were well-aligned on the pseudotime trajectory, and clustered close in both the early (quiescent) and late (differentiation) stages of pseudotime (**Fig 4D**, sham and day 7). In contrast, the MuSCs/MPCs exhibited drastic strain-specific differences at 1dpi and 3dpi. At both of these times, a large fraction of C57BL/6 MuSCs/MPCs were in the activation/proliferation phase (**Fig 4D**, days 1 and 3, blue curve), whereas the vast majority of the BALB/c MuSCs/MPCs occupied the late, differentiation stage of the pseudotime trajectory. This pattern of MuSC/MPC distribution along the pseudotime trajectory suggests that the BALB/c MPCs committed to differentiation prematurely. This conclusion is also supported by immunohistochemical analysis, which revealed that at 7dpi BALB/c TA muscle contained terminally differentiated eMHC+ new myofibers, while the numbers of new muscle fibers and Pax7+ MuSCs were much less compared to C57BL/6 muscle (**Fig 2C, 2D**). Taken together, these results suggest that the failure of skeletal muscle regeneration in the BALB/c model of CLTI is at least partially due to the premature differentiation of MuSCs/MPCs.

### The pro-inflammatory niche induces premature differentiation of MuSCs in BALB/c mice following HLI

To address whether the inflammatory macrophages in BALB/c muscle are associated with premature differentiation of MuSCs following HLI, we first used RNAscope to assess the spatial distribution of F4/*80*+ (macrophage marker) and *Myod1*+ (MuSC/MPC marker) cells in TA muscle at 3dpi of HLI. This approach demonstrated the proximity of macrophages to MuSCs in both C57BL/6 and BALB/c mice in the ischemic limb (**Fig 4E**). Next, we assessed the probability of intercellular communication between macrophages and MuSCs using a computational method called CellphoneDB ^37^. We found that at 3dpi, C57BL/6 MuSCs are likely responsive to well-characterized pro-regenerative cytokines secreted by macrophages, including TGFB1 and IGF1 **(Fig 4F)**, which are known to play critical roles in regulating skeletal muscle regeneration by promoting MPC proliferation and preventing MPC differentiation both in vitro and in vivo ^33,38,39^. In contrast, a role for TGFB1 and IGF1 in macrophage-MuSC intercellular communication was not detected in BALB/c mice. Moreover, the genes highly expressed in C57BL/6 MuSCs at 3dpi, including *Igf1r* (**Fig S4C**), were significantly enriched for GO terms associated with skeletal muscle cell proliferation (**Fig 4G**). These results suggest that the aberrant inflammatory state of macrophages disrupts the pro-regenerative signaling communication between macrophages and MuSCs, such as IGF1, which is essential for MuSC expansion and the prevention of premature differentiation. Dysregulation of macrophage-MuSC signaling contributes to the premature differentiation phenotype observed in MuSCs of BALB/c mice.

Finally, to determine the causal role of macrophage-secreted ligands in regulating MuSC/MPC proliferation and differentiation, we purified primary MuSCs from both mouse strains for in vitro cell proliferation assays using BrdU incorporation. Upon treatment with recombinant IGF1, new DNA synthesis in MuSCs/MPCs isolated from both mouse strains significantly increased by 2-3 fold (**Fig 4H**). Proliferation and differentiation of MuSCs/MPCs exhibit a remarkable inverse relationship, as the myogenic differentiation program necessitates cell-cycle exit and inhibition of proliferative machinery. Consequently, we reasoned that in BALB/c muscle, the lack of expression of pro-regenerative cytokines, such as IGF1, from inflamed macrophages at least partially contributes to the premature differentiation of MuSCs/MPCs and failure of limb muscle regeneration in response to ischemia.

### Disruption of MuSCs fate switch is associated with aberrant macrophage-MuSC signaling crosstalk in human CLTI

Finally, we sought to translate our findings from murine models of CLTI to CLTI patients. Because of the inverse relationship between proliferation and differentiation, we reasoned that if premature differentiation of MuSCs also occurs in human CLTI, it would lead to reduced MuSC/MPC numbers in ischemia-injured distal limbs. Consistent with this possibility, we found that the distal muscle of two out of three CLTI patient samples analyzed by scRNA-seq contained ∼60% fewer MuSCs/MPCs compared to matched proximal muscle (**Fig S5A**). To further explore these findings, we conducted Pax7 immunostaining on cross-sections of limb muscles collected from another seven CLTI patients (**Fig S5B**, representative images from one patient). In four out of the seven patients, the numbers of Pax7+ MuSCs in the ischemia-injured distal muscle were ∼10-60% less than those in the patient matched non-ischemic proximal muscle (**Fig S5B, S5C**). These results indicate that in at least a subset of CLTI patients, approximately 57% - 67%, ischemia-injured limb muscles contain fewer MuSCs/MPCs compared to non-ischemic muscle.

To assess changes in gene expression and signaling activity of MuSCs/MPCs in human CLTI, we performed an integrative analysis of our human CLTI scRNA-seq data in conjunction with published human muscle scRNA-seq data generated from 10 healthy individuals ^40^ (**Fig 5A, S5D**). This analysis revealed Pax7+ MuSCs, MyoG+ MPCs populations in human muscle, and the C3AR1+ macrophage population (**Fig 5B, S5E - S5G**). Notably, we inferred intercellular communications ^41^ using scRNA-seq data and discovered that both the number and strength of signaling pathways between MuSCs/MPCs and macrophages were substantially stronger in the distal, ischemic muscle than in the proximal, non-ischemic human muscle (**Fig 5C**). The top-ranked inter-cellular signaling flow from macrophages to MuSCs/MPCs in the distal tissue included well-established pro-inflammatory pathways, such as SPP1, CCL, TNF, and CXCL (**Fig 5D**), highlighting the aberrant signaling communication between inflammatory macrophages and MuSCs/MPCs/myoblasts in ischemia-injured human CLTI limb muscle. Moreover, we separated quiescent MuSCs that were Pax7 high (**Fig 5E, 5F, S5F** clusters 0 and 1) from MPCs committed to myogenic differentiation and expressing MYOG/MYH2 mRNA (**Fig 5E, 5F, S5F** clusters 2 and 4) on a new UMAP space. Significant pro-inflammatory signaling between macrophages and MuSCs/MPCs, such as IL6-IL6R, CCL4-SLC7A1, and SPP1-CD44/PTGER4, was detected only in the ischemic distal tissue and not in the non-ischemic proximal tissue (**Fig 5G**). Notably, distal MuSCs/MPCs expressed significantly lower levels of the MuSC quiescence/self-renewal marker SPRY1 and higher levels of the differentiation marker MYOG compared to the proximal tissue of CLTI patients and healthy individuals (**Fig 5H**). These findings collectively demonstrate that the permanent tissue loss phenotype of human CLTI is associated with increased pro-inflammatory signaling between inflammatory macrophages and MuSCs and MuSC-derived myogenic cells. Importantly, the ischemia-damaged MuSCs and myoblasts exhibit gene expression signatures indicating a loss of quiescence (lower SPRY1) and premature differentiation (higher MYOG) compared to the non-ischemic tissue of CLTI patients and healthy muscle.

**Figure 5.**
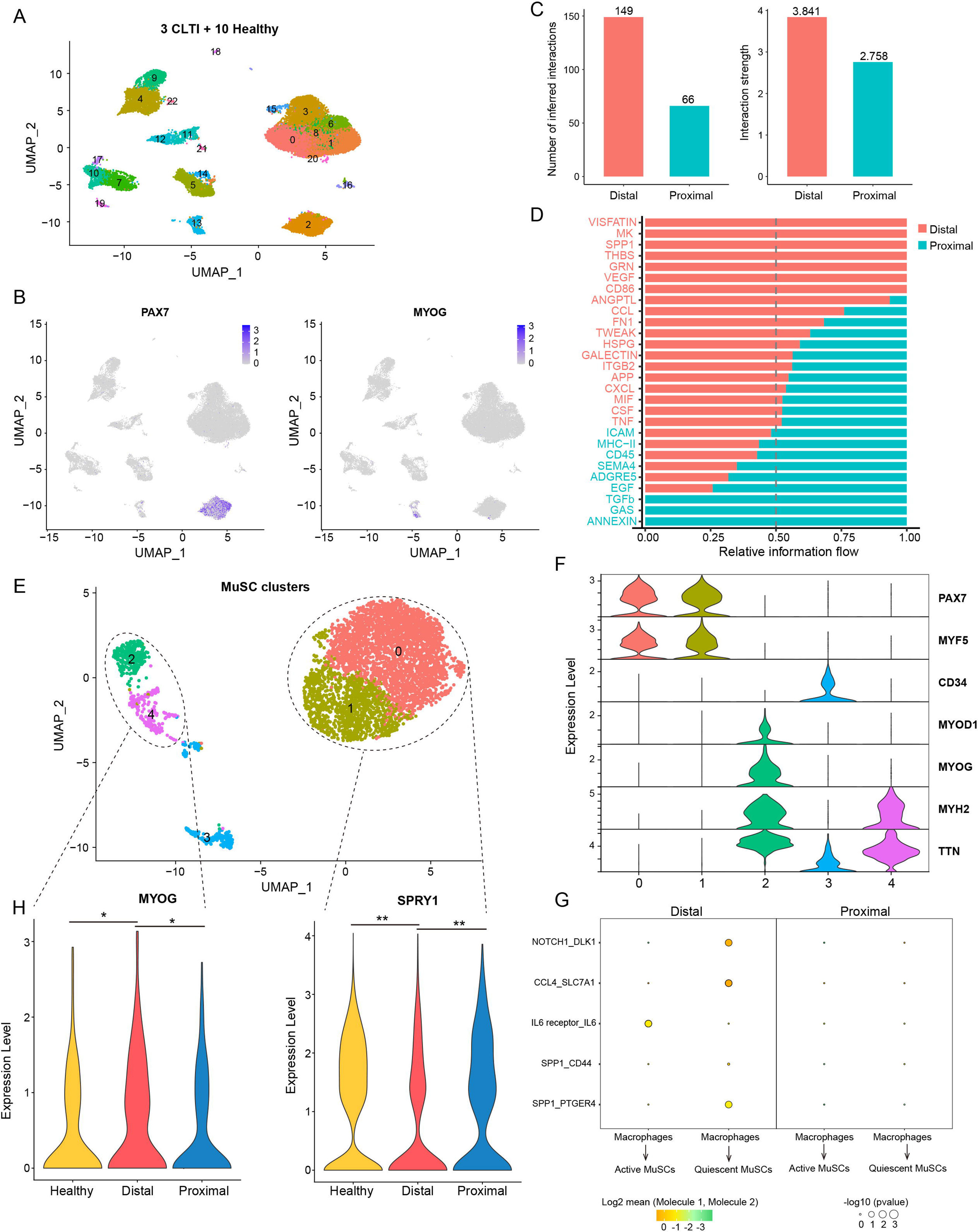
Increased pro-inflammatory signaling between macrophages and MuSC/MPCs and dysregulated cell state of MuSC/MPCs in the ischemia-damaged human limb of CLTI patients. **A:** UMAP visualization of single-cell data from human skeletal muscle in 3 CLTI patients, including both proximal and distal conditions, and 10 healthy individuals. A total of 34,950 cells are plotted on the UMAP. **B:** Feature plots displaying gene expression of PAX7 mRNA in quiescent MuSCs and MyoG mRNA in activated MuSC/MPCs. **C:** The inferred strength and the number of signaling interactions calculated by CellChat between macrophages and MuSCs in ischemic and non-ischemic muscles of CLTI patients. **D:** The significant signaling pathways between macrophages and MuSCs/MPCs are ranked based on their inferred strength differences between ischemic and non-ischemic skeletal muscles. Signaling pathways colored in red are enriched in ischemic muscle, while those colored in blue are enriched in non-ischemic conditions. **E, F: (E)** The MuSC/MPCs (clusters 2 and 13 shown in **A**) are plotted on a new UMAP space with increased resolution to visualize the differences in MuSCs/MPCs. **(F)** In this new UMAP, cells in clusters 2 and 4, defined as active MuSCs, express high levels of differentiation marker MyoG, while cells in clusters 0 and 1, defined as quiescent MuSCs, express high levels of quiescent marker Pax7. **G:** Ligand-receptor interactions inferred by CellPhoneDB between macrophages and activated/quiescent MuSCs in ischemic and non-ischemic conditions. In distal conditions, we observed stronger pro-inflammatory signaling pathways, such as IL6, CCL4, and SPP1, compared to proximal conditions. **H:** In distal muscle of CLTI patients, the active MuSCs express the highest level of myogenic differentiation marker MyoG, indicating precocious differentiation; while quiescent MuSCs express the lowest level of quiescent/self-renewal marker SPRY1, indicating loss of stem cell quiescence. Violin plots show the expression of MyoG and SPR1 in active and quiescent MuSCs in the indicated conditions in healthy and CLTI patients. * P-value < 0.05; ** P-value < 0.01. P-values were calculated using the FindMarkers function of the Seurat R package with the Wilcoxon test method.

## DISCUSSION

In this study, we utilized a rigorous experimental approach with the acquisition of ischemic and non-ischemic macrophages from the same patient for our analysis. This approach limits the influence of confounding variables, such as diabetes, smoking and age, which are ubiquitous in human studies. Through analyses of human CLTI scRNA-seq datasets, our work provides evidence suggesting that the pro-inflammatory macrophages are associated with and at least partially contribute to the permanent tissue loss phenotype in CLTI. Moreover, these findings are further validated in the murine models of CLTI. A recent histologic assessment of macrophages and MuSCs obtained from gastrocnemius muscle biopsies in human patients demonstrated that anti-inflammatory (CD206+) macrophages were associated with increased MuSC content and muscle fiber size ^30^. Taking all these results into consideration, our findings suggest a critical role of macrophage and inflammation in determining CLTI progression.

Furthermore, our findings here indicate that the macrophages in the ischemic damaged tissue create an inflamed niche that disrupts muscle regeneration by inducing precocious differentiation of MuSCs. Identifying the paracrine factors produced by the macrophages that mediate this effect may therefore throw light on potential therapeutic interventions. We have presented evidence for IGF1 signaling pathways which are required for muscle regeneration but lost in the ischemic damaged limbs. Moreover, our analysis revealed additional pathways that are differentially activated between macrophages and MuSCs in the BALB/c and C57BL/6 strains. For example, we find that SPP1-CD44 signaling is highly active in macrophage-MuSC ligand-receptor pairs in both a murine CLTI model (BALB/c) and human CLTI patients. SPP1 is a pro-inflammatory cytokine that inhibits MuSC proliferation ^42^. Hence, SPP1 signaling represents a potential mechanism to support our findings that the ischemic damaged MuSCs proceed to premature myogenic. In the future study, functional and mechanistic investigation of these pathways in both engineered human muscle bundles ^43^ and the murine models of CLTI may shed light to develop therapeutic strategies to manipulate MuSCs and improve tissue repair.

In summary, our work represents a significant advancement in our collective understanding of macrophage in the ischemic human limb at the cellular, transcriptomic, and signaling levels. Our findings point to new cellular mechanisms that can be potentially exploited to improve muscle function and lay a foundation for future muscle-specific therapies for limb salvage.

### Accession Codes and Data Availability

Sequencing data has been deposited in the NCBI Gene Expression Omnibus (GEO) (http://www.ncbi.nlm.nih.gov/geo) under accession number GSE227077. The embargo will be lifted upon manuscript acceptance for publication. Additional materials, data, code, and related protocols are available upon request.

### Author contributions

K.W.S., Y.Xu, D.T.P. and Y.D. designed the study. Y.Xu led bioinformatics data analysis with input from Y.Xiang. D.T.P. generated the experimental data with help from X.W., X.L., K.F., L.A.O., J.M.M., J.O., Q.D.. K.W.S., C.D.K. and Y.D. supervised the study. K.W.S., Y.Xu., D.T.P., K.F. and Y.D. wrote the paper.

## Acknowledgments

We thank Drs. Brigid Hogan (Duke) for feedback on previous versions of the manuscript. This work is supported by a new lab startup fund from Duke University Regeneration Next Initiative (current Duke Regeneration Center) (to Y.D.), Duke Whitehead Scholarship (to Y.D.), Glenn Foundation for Medical Research and AFAR Grants for Junior Faculty (to Y.D.), NIH 4D Nucleome Consortium U01HL156064 (to Y.D.), and NIH Genomic Innovator Awards R35HG011328 (to Y.D.). Duke University Medical Center Physician-Scientist Strong Start Award (to K.W.S.). Y. Xiang is supported by a Duke Regeneration Center postdoctoral fellowship. Y. Xu is supported by the Center for Advanced Genomic Technologies postdoctoral fellowship and the Duke Regeneration Center Fellowship to Accelerate Career Independence.

## MATERIALS AND METHODS

### PCR primers and oligo nucleotides used in this study are listed in Table S4

### Human tissue collection and single-cell isolation

Skeletal muscle was obtained from CLTI patients (*n* = 3) undergoing lower-limb amputation in accordance with a research protocol approved by the Duke University Institutional Review Board (IRB#Pro00065709). Paired samples from proximal and distal muscle bodies were collected and subject to mechanical dissociation. A subsequent enzymatic digestion was performed using either 0.05% pronase (Sigma, 537088) for 1 hour (Patient #1) or 3.7mg/mL collagenase II (Worthington, LS004177) for 90 minutes followed by 6mg/mL dispase (Gibco, 17105-041) for 30 minutes (Patients #2 and #3). Cells were passed through a 100uM Steriflip vacuum filter (EMD Millipore, SCNY00100) and resuspended in Ham’s F-10 media supplemented with 10% horse serum and 1x penicillin/streptomycin. The single cell suspensions were stained with Propidium Iodide (PI). Fluorescence-activated cell sorting (FACS) was performed using a SonySorter SH800S to isolate PI-live cells. For human samples, approximately 150,000 PI-live cells were sorted per sample and 16,000-24,000 cells used for scRNA-seq library generation.

### Mouse hind limb ischemic injury and single-cell isolation

Hind limb ischemia (HLI) surgery was performed on C57BL/6 and BALBc mice. To induce muscle ischemia, the femoral artery was ligated proximally, inferior to the inguinal ligament just proximal to the lateral circumflex femoral artery, as well as distally, immediately proximal to the bifurcation of the popliteal and saphenous arteries ^44^. Laser Doppler Perfusion Imaging (LDPI) was performed with a Moor Instruments LDI2-High Resolution (830nM) System (Moor, Axminster, UK) to quantify and assess blood flow restoration.

### Mouse tissue collection and single-cell isolation

Mouse hindlimb muscles were collected on days 0 (no injury), 1, 3, and 7 following HLI surgery and single-cell suspensions were generated using mechanical dissociation followed by enzymatic digestion with 0.05% Pronase (Sigma, 537088) for one hour. Cells were vacuum filtered as described above and subsequently stained with PI, anti-CD45-Alexa Fluor 488 (clone HI30, Invitrogen, MHCD4520), and FITC anti-CD31 (BioLegend, 102506). Using FACS, 150,000 PI-cells and an additional 150,000 PI-/CD31-/CD45-cells were isolated. The latter population (“depleted” cells) was isolated to increase the representation of non-hematopoietic and non-endothelial cells, given the relative scarcity of muscle stem cells and fibro-adipogenic progenitors. Live cells and “depleted” cells were pooled at a 1:1 ratio and a total of 16,000-24,000 cells used for scRNA sequencing.

### Single-cell RNA sequencing library generation

Cells were collected for single-cell RNA-seq analysis as described above. Single-cell RNA-seq libraries were generated using Chromium Next GEM Single Cell 3’ Reagent Kits v3.1 (10x Genomics) according to the manufacturer’s protocol.

### Immunofluorescent staining

Human skeletal muscle samples from both ischemic (distal) and non-ischemic (proximal) muscle were obtained from surgical amputation specimens. Tissue was harvested and embedded in OCT compound using liquid nitrogen. 8 μm sections were prepared on microscope slides using cryostat sectioning for histological analysis. Frozen sections were allowed to come to room temperature and fixed in 4% paraformaldehyde for 10 minutes, permeabilized with 0.1% Triton X-100 in PBS for 5 minutes and then washed in PBS. Blocking solution (5% normal donkey serum in PBS) was applied for 30 minutes at room temperature followed by primary antibody staining overnight at 4°C using anti-CD206 (R&D systems, AF2534), anti-CD11b (Cell Sciences, MON1019-1, clone Bear-1), and anti-dystrophin (Thermo Scientific, RB-9024-P). Tissue sections were then washed using PBS, incubated with the secondary antibody (ThermoFisher Scientific) for 1 hour at room temperature, and counterstained with 1ug/mL Hoechst 33342 (Thermo Scientific, 6629) for 5 minutes at room temperature. The tissue sections were mounted with Fluoromount-G™ Mounting Medium (ThermoFischer Scientific, 00-4958-02). Images were acquired using a Zeiss Axio Imager Z2 Upright Microscope at x200 magnification.

### Macrophage isolation using FACS

Hind limb muscles were collected from C57BL/6 and BALB/c mice at day 3 after HLI surgery. Single cell suspensions were generated as described above. Cells were blocked with purified anti-mouse CD16/32 antibody (Biolegend, #101301) for 10 minutes. Primary antibody staining was performed using anti-CD45-Alexa Fluor 488 (clone HI30, Invitrogen, MHCD4520), anti-CD11b (clone M1/70, Invitrogen, 12-0112-81), and anti-F4/80-biotin (clone A3-1, Bio-rad, MCA497BT) antibodies for 40 minutes. Streptavidin-PE/Cy7 (Biolegend, 405206) was used as a secondary reagent for anti-F4/80-biotin (PMID: 25896247). Using FACS, macrophages were isolated by PI-/CD45+/CD11b+/F480+ gating.

### Bulk RNA sequencing

Total RNA was extracted using TRIzol Reagent (Invitrogen) according to manufacturer’s protocol. First-strand reverse transcription and template switching was performed using an Oligo(dT) primer (dT30VN-ME-A), a locked nucleic acid-containing TSO (NotI-TSO), and Superscript IV reverse transcriptase (Invitrogen, # 18090050). PCR preamplification of cDNA was performed with IS PCR and Tn5ME-A-aHic using 2X KAPA PCR mix (Kapa Biosystem, KK2602) followed by cleanup using SPRISelect beads (Beckman Coulter, REF B23319). DNA was digested by NotI-HF (NEB, #R3189L), tagmented using Tn5 assembled with adaptors Tn5ME-A/Tn5MErev and Tn5ME-B/Tn5MErev. A Zymo DNA clean and concentrator kit (Zymo, R1014) was utilized ^45^. Library PCR was performed using unique combinations of Nextera-PCR i5/i7 primers. DNA strands between 400-600bp were selected by gel extraction using Zymoclean Gel DNA recovery kit (Zymo, #D4002). DNA libraries were submitted for next-generation sequencing using paired-end sequencing.

### RNAscope *In situ* hybridization procedure

Mouse tibialis anterior was harvested on day 3 after HLI and embedded in OCT compound in ice-cold isopentane. Cryostat sections (10 μm) were prepared on microscope slides (Superfrost Plus, Fisher Scientific), and immediately stored at −80°C until use. Samples for RNAscope® (Advanced Cell Diagnostics) were processed according to the manufacturer’s guidelines for preparing fresh frozen tissue. Muscle sections were pre-treated with protease IV (RNAscope® protease III and IV reagents, ACD, 322340) for 15 minutes and incubated in the desired probes [ADGRE1 (F4/80), ACD) and Myod1 (ACD] for 2 hours at 40°C in the EZ Hybridization oven. RNAscope *In situ* hybridization was performed using RNAscope® Fluorescent Multiplex Assay Detecting Reagents kit (ACD, 320851) according to the manufacturer’s protocols. Images were acquired using a Zeiss Axio Imager Z2 Upright Microscope at x400 magnification. Images analysis was performed according to the RNAscope® Data Analysis guide (ACD).

### Cell proliferation assay

Skeletal muscle stem cells were isolated from uninjured young adult C57BL/6 and BALB/c hindlimb skeletal muscle and myogenic cells were expanded in culture using an established protocol ^46^. Cells were seeded in 96-well plates and 24 hours after plating treated with: THBS1 (R&D systems, 7859-TH-050), syndecan-4 Ab (BD Pharmingen, 550350), normal rat IgG2a (EMD Millipore, MABF 1077Z), IGF-1 (R&D systems, 791-MG), FGF2 (Thermo Fisher, PHG0367) or vehicle (PBS) for 24 hours. After 72 hours of culture with the indicated ligands EdU was added to the medium, the cells were cultured for an additional 6 hours, and then fixed. Cells were analyzed using the Click-iT EdU Cell Proliferation Kit for Imaging (Invitrogen, C10337) according to the manufacturer’s instructions.

### Single cell RNA-seq data processing and analysis

The sequencing reads (10× Genomics) were processed using the Cell Ranger pipeline (v3.1.0,) with GRCm38 reference genome. The output filtered matrices of different samples as input files for downstream analyses using the R package Seurat (v4.0.1) ^47^. Genes expressed by less than 3 cells were removed. Cells with unique feature counts over 4100, under 1000, or greater than 25% mitochondrial RNA counts were filtered. Meanwhile, cells that were recognized as doublets by the Python package Scrublet (v0.2.3) were removed. The different Seurat objects from different samples were combined to create a new object.

The gene expression levels of the combined object for each cell were normalized and log-transformed by NormalizeData function. Scaled data of all cells by ScaleData function was used for principal component analysis based on 2000 highly variable genes with RunPCA function. The harmony algorithm was then used to correct the batch correction ^48^. The resulting Harmony embeddings instead of PCA was used in non-linear dimensional reduction and nearest neighbor graph construction. The clusters were defined using FindClusters function and marker genes for each cluster were identified by running the FindAllMarkers function. Cell type of the clusters were annotated according to these marker genes. These steps were also used to human scRNA-seq data analysis and mouse macrophage sub-clusters. The AddModuleScore function was used to calculate module scores for feature expression programs related to inflammatory response in macrophages. The differentially expressed genes (DEGs) were identified by FindMarkers function. The gene sets for the macrophage GSEA analysis were download from MSigDB. ^49–51^

The satellite cells were exacted from the Seurat object of each sample for the sctransform normalization and then were merged as a new object. The SelectIntegrationFeatures function was used to choose the top scoring features for the satellite cell integrated object. Then the umap and sub-clusters were defined with the above steps (RunPCA, RunHarmony, RunUMAP, FindNeighbors, FindClusters). The trajectory inference and pseudotime calculations on the integrated object were performed using the Monocle3 ^52^. Cell-cell communication between macrophages and satellite cells was inferred using CellphoneDB v3.1 ^37,53^ and CellChat v1.4^41^.

### Bulk RNA-seq data analysis

The reads from bulk RNA-seq library were aligned to GRCm38 reference genome by STAR v2.7.4a with “--sjdbOverhang 99” ^54^. FeatureCounts v1.6.3 was used to determine the read counts for each gene ^55^. The DEGs were identified using DESeq2 v1.30.1 with the threshold that adjusted p-value less than_J0.05 and fold change greater than 2. GO enrichment analysis was performed with these DEGs via DAVID ^56,57^. The bigwig files generated by bamCoverage v3.5.1 ^58^ with RPKM normalization method from alignment of reads (bam files) were used to read coverage visualization through Integrative Genomics Viewer (IGV v 2.11.9)^59^

## SUPPLEMENTARY FIGURES AND TABLES

**Supplementary Figures 1-6, and Supplementary Tables 1-4.**

**Supplementary Figure 1. Single-cell transcriptome analysis of skeletal muscle in human CLTI patients. Related to Figure 1.**

**A:** Representative computed tomography angiography images from three PAD patients, demonstrating similar atherosclerotic lesions and degrees of perfusion in patients with disparate clinical phenotypes. Ankle brachial index (ABI).

**B:** UMAP embedding of single-cell RNA-seq profiles from non-ischemic and ischemic skeletal muscle samples from three CLTI patients. Cells are colored by patient and tissue condition.

**C:** Feature plots showing the expression of the indicated cell type-specific marker genes.

**D:** Representative immunofluorescence images of macrophages in ischemic and non-ischemic muscle of CLTI patients. CD11b+ (green), CD206+ (red). DAPI (blue).

**Supplementary Figure 2. Single-cell RNA-seq atlas of limb muscle regeneration and damage in C57BL/6 and BALB/c mouse strains following HLI surgery. Related to Figure 2.**

**A:** UMAP visualization of cells from BALB/c mice at indicated times before and after HLI surgery.

**B:** UMAP visualization of cells from C57BL/6 mice at indicated times before and after HLI surgery.

**C:** Feature plots highlighting mRNA expression of the indicated cell type-specific marker genes in each cell cluster.

**Supplementary Figure 3. Distinct macrophage populations in C57BL/6 and BALB/c mice following limb ischemia. Related to Figure 3.**

A: UMAP visualization of all macrophages from two mouse strains and four time points. Total number of macrophages: n=26,991.

**B:** Heatmap showing expression levels of marker genes in each sub-cluster of macrophages shown in **A**.

**C:** FACS plot showing that CD45+/CD11b+/F4/80+ macrophages were purified from the hindlimb muscle of the two mouse strains at 3 days after HLI for bulk RNA-seq analysis.

**D:** Genome browser views of bulk RNA-seq data at Gdf3 and Igf1 loci in macrophages purified from the two mouse strains at day 3 post-HLI surgery.

**Supplementary Figure 4. Single-cell analysis of MuSCs/MPCs in C57BL/6 and BALB/c mice before and after HLI surgery. Related to Figure 4.**

**A:** Feature plots highlighting mRNA expression of the indicated genes in sub-clusters of MuSCs/MPCs.

**B:** UMAP embedding of the pseudotime trajectories of MuSCs/MPCs. Color key: pseudotimes.

**C:** Violin plot showing Igf1r expression in MuSC/MPCs of BALB/c versus C57BL/6 mice at 3 days post-HLI surgery.

**Supplementary Figure 5. Dysregulation of macrophage-MuSC crosstalk and MuSC cell states in ischemia-damaged human muscle of CLTI patients. Related to Figure 5.**

**A:** Proportion of MuSCs/MPCs among all cells in proximal and distal skeletal muscle tissue from three CLTI patients. Cell numbers are quantified by scRNA-seq data.

**B:** Representative immunofluorescence images of MuSCs (PAX7+) in ischemic (distal, left) and non-ischemic (proximal, right) skeletal muscle specimens from a CLTI patient. PAX7+ (red). Laminin (green). Hoechst (blue). CLTI patient number: N=7.

**C:** Quantification of (B) using data from 7 representative CLTI patients.

**D:** UMAP visualization of cells from skeletal muscle tissues of 10 healthy individuals and 3 CLTI patients. Color key indicates the patient and condition (proximal versus distal). Total number of cells: n=34,950.

**E:** Feature plot showing the expression of human macrophage marker C3AR1 in cluster 7 shown in Fig. 5A.

**F:** The MuSCs/MPCs were plotted on a new UMAP space with increased resolution. Feature plots highlight the expression of the indicated genes in sub-clusters of MuSCs/MPCs.

**G:** Distribution of the MuSCs/MPCs obtained from the three CLTI patients in both distal (left) and proximal (right) conditions.

**Table S1: The differentially expressed genes in macrophage clusters 1 and 2 versus cluster 0 shown in Figure 1E. Gene expression is quantified by scRNA-seq data. Related to Figure 1.**

**Table S2: The top 20 marker genes for each sub-cluster of macrophages shown in Figure S3A and S3B. Related to Figure 3 and Figure S3.**

**Table S3: The differentially expressed genes in macrophage purified from BALB/c versus C57BL6 mice at 3 days post HLI surgery. Gene expression is quantified by bulk RNA-seq. Related to Figure 3**.

**Table 4. List of PCR primers used in this study.**

**Table S4.**
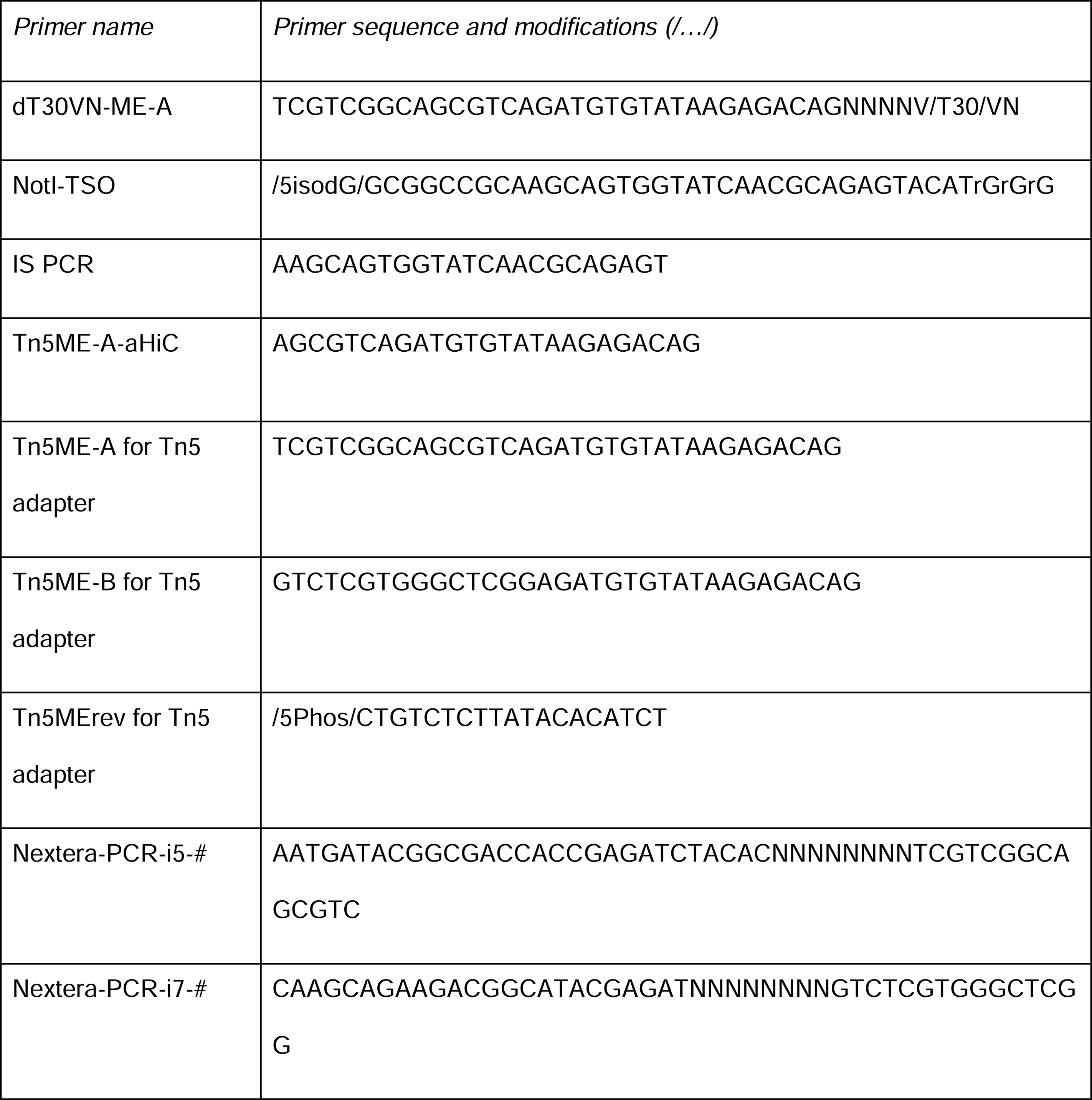
List of PCR primers used in this study.

## REFERENCES

1. Gutierrez, J.A., Aday, A.W., Patel, M.R., and Jones, W.S. (2019). Polyvascular Disease: Reappraisal of the Current Clinical Landscape. Circ. Cardiovasc. Interv. 12, e007385.

2. Morley, R.L., Sharma, A., Horsch, A.D., and Hinchliffe, R.J. (2018). Peripheral artery disease. BMJ 360, j5842.

3. Kullo, I.J., and Rooke, T.W. (2016). CLINICAL PRACTICE. Peripheral Artery Disease. N. Engl. J. Med. 374, 861–871.

4. Fowkes, F.G.R., Aboyans, V., Fowkes, F.J.I., McDermott, M.M., Sampson, U.K.A., and Criqui, M.H. (2017). Peripheral artery disease: epidemiology and global perspectives. Nat. Rev. Cardiol. 14, 156–170.

5. Criqui, M.H., and Aboyans, V. (2015). Epidemiology of peripheral artery disease. Circ. Res. 116, 1509–1526.

6. Farber, A. (2018). Chronic Limb-Threatening Ischemia. N. Engl. J. Med. 379, 171–180.

7. Duff, S., Mafilios, M.S., Bhounsule, P., and Hasegawa, J.T. (2019). The burden of critical limb ischemia: a review of recent literature. Vasc. Health Risk Manag. 15, 187–208.

8. Farber, A., Menard, M.T., Conte, M.S., Kaufman, J.A., Powell, R.J., Choudhry, N.K., Hamza, T.H., Assmann, S.F., Creager, M.A., Cziraky, M.J., et al. (2022). Surgery or Endovascular Therapy for Chronic Limb-Threatening Ischemia. N. Engl. J. Med. 387, 2305–2316.

9. Fadini, G.P., Spinetti, G., Santopaolo, M., and Madeddu, P. (2020). Impaired Regeneration Contributes to Poor Outcomes in Diabetic Peripheral Artery Disease. Arterioscler. Thromb. Vasc. Biol. 40, 34–44.

10. Aranguren, X.L., Verfaillie, C.M., and Luttun, A. (2009). Emerging hurdles in stem cell therapy for peripheral vascular disease. J. Mol. Med. 87, 3–16.

11. Pizzimenti, M., Meyer, A., Charles, A.-L., Giannini, M., Chakfé, N., Lejay, A., and Geny, B. (2020). Sarcopenia and peripheral arterial disease: a systematic review. J. Cachexia Sarcopenia Muscle 11, 866–886.

12. McClung, J.M., McCord, T.J., Ryan, T.E., Schmidt, C.A., Green, T.D., Southerland, K.W., Reinardy, J.L., Mueller, S.B., Venkatraman, T.N., Lascola, C.D., et al. (2017). BAG3 (Bcl-2-Associated Athanogene-3) Coding Variant in Mice Determines Susceptibility to Ischemic Limb Muscle Myopathy by Directing Autophagy. Circulation 136, 281–296.

13. McClung, J.M., McCord, T.J., Southerland, K., Schmidt, C.A., Padgett, M.E., Ryan, T.E., and Kontos, C.D. (2016). Subacute limb ischemia induces skeletal muscle injury in genetically susceptible mice independent of vascular density. J. Vasc. Surg. 64, 1101–1111.e2.

14. Polonsky, T.S., and McDermott, M.M. (2021). Lower Extremity Peripheral Artery Disease Without Chronic Limb-Threatening Ischemia: A Review. JAMA 325, 2188–2198.

15. Conte, M.S. (2017). Data, guidelines, and practice of revascularization for claudication. J. Vasc. Surg. 66, 911–915.

16. Sukul, D., Grey, S.F., Henke, P.K., Gurm, H.S., and Grossman, P.M. (2017). Heterogeneity of Ankle-Brachial Indices in Patients Undergoing Revascularization for Critical Limb Ischemia. JACC Cardiovasc. Interv. 10, 2307–2316.

17. Cong, G., Cui, X., Ferrari, R., Pipinos, I.I., Casale, G.P., Chattopadhyay, A., and Sachdev, U. (2020). Fibrosis Distinguishes Critical Limb Ischemia Patients from Claudicants in a Transcriptomic and Histologic Analysis. J. Clin. Med. Res. 9. 10.3390/jcm9123974.

18. Ryan, T.E., Yamaguchi, D.J., Schmidt, C.A., Zeczycki, T.N., Shaikh, S.R., Brophy, P., Green, T.D., Tarpey, M.D., Karnekar, R., Goldberg, E.J., et al. (2018). Extensive skeletal muscle cell mitochondriopathy distinguishes critical limb ischemia patients from claudicants. JCI Insight 3. 10.1172/jci.insight.123235.

19. McClung, J.M., McCord, T.J., Keum, S., Johnson, S., Annex, B.H., Marchuk, D.A., and Kontos, C.D. (2012). Skeletal muscle-specific genetic determinants contribute to the differential strain-dependent effects of hindlimb ischemia in mice. Am. J. Pathol. 180, 2156–2169.

20. Sousa-Victor, P., García-Prat, L., and Muñoz-Cánoves, P. (2022). Control of satellite cell function in muscle regeneration and its disruption in ageing. Nat. Rev. Mol. Cell Biol. 23, 204–226.

21. van Velthoven, C.T.J., and Rando, T.A. (2019). Stem Cell Quiescence: Dynamism, Restraint, and Cellular Idling. Cell Stem Cell 24, 213–225.

22. Chazaud, B. (2020). Inflammation and Skeletal Muscle Regeneration: Leave It to the Macrophages! Trends Immunol. 41, 481–492.

23. Dort, J., Fabre, P., Molina, T., and Dumont, N.A. (2019). Macrophages Are Key Regulators of Stem Cells during Skeletal Muscle Regeneration and Diseases. Stem Cells Int. 2019, 4761427.

24. Rybalko, V., Hsieh, P.-L., Merscham-Banda, M., Suggs, L.J., and Farrar, R.P. (2015). The Development of Macrophage-Mediated Cell Therapy to Improve Skeletal Muscle Function after Injury. PLoS One 10, e0145550.

25. Wosczyna, M.N., and Rando, T.A. (2018). A Muscle Stem Cell Support Group: Coordinated Cellular Responses in Muscle Regeneration. Dev. Cell 46, 135–143.

26. Tidball, J.G. (2017). Regulation of muscle growth and regeneration by the immune system. Nat. Rev. Immunol. 17, 165–178.

27. Varga, T., Mounier, R., Horvath, A., Cuvellier, S., Dumont, F., Poliska, S., Ardjoune, H., Juban, G., Nagy, L., and Chazaud, B. (2016). Highly Dynamic Transcriptional Signature of Distinct Macrophage Subsets during Sterile Inflammation, Resolution, and Tissue Repair. J. Immunol. 196, 4771–4782.

28. Shi, C., and Pamer, E.G. (2011). Monocyte recruitment during infection and inflammation. Nat. Rev. Immunol. 11, 762–774.

29. Kosmac, K., Peck, B.D., Walton, R.G., Mula, J., Kern, P.A., Bamman, M.M., Dennis, R.A., Jacobs, C.A., Lattermann, C., Johnson, D.L., et al. (2018). Immunohistochemical Identification of Human Skeletal Muscle Macrophages. Bio Protoc 8. 10.21769/BioProtoc.2883.

30. Kosmac, K., Gonzalez-Freire, M., McDermott, M.M., White, S.H., Walton, R.G., Sufit, R.L., Tian, L., Li, L., Kibbe, M.R., Criqui, M.H., et al. (2020). Correlations of Calf Muscle Macrophage Content With Muscle Properties and Walking Performance in Peripheral Artery Disease. J. Am. Heart Assoc. 9, e015929.

31. Schmidt, C.A., Amorese, A.J., Ryan, T.E., Goldberg, E.J., Tarpey, M.D., Green, T.D., Karnekar, R.R., Yamaguchi, D.J., Spangenburg, E.E., and McClung, J.M. (2018). Strain-Dependent Variation in Acute Ischemic Muscle Injury. Am. J. Pathol. 188, 1246–1262.

32. Gordon, S. (2003). Alternative activation of macrophages. Nat. Rev. Immunol. 3, 23–35.

33. Tonkin, J., Temmerman, L., Sampson, R.D., Gallego-Colon, E., Barberi, L., Bilbao, D., Schneider, M.D., Musarò, A., and Rosenthal, N. (2015). Monocyte/Macrophage-derived IGF-1 Orchestrates Murine Skeletal Muscle Regeneration and Modulates Autocrine Polarization. Mol. Ther. 23, 1189– 1200.

34. Patsalos, A., Simandi, Z., Hays, T.T., Peloquin, M., Hajian, M., Restrepo, I., Coen, P.M., Russell, A.J., and Nagy, L. (2018). In vivo GDF3 administration abrogates aging related muscle regeneration delay following acute sterile injury. Aging Cell 17, e12815.

35. Lepper, C., Partridge, T.A., and Fan, C.-M. (2011). An absolute requirement for Pax7-positive satellite cells in acute injury-induced skeletal muscle regeneration. Development 138, 3639–3646.

36. Relaix, F., Bencze, M., Borok, M.J., Der Vartanian, A., Gattazzo, F., Mademtzoglou, D., Perez-Diaz, S., Prola, A., Reyes-Fernandez, P.C., Rotini, A., et al. (2021). Perspectives on skeletal muscle stem cells. Nat. Commun. 12, 692.

37. Efremova, M., Vento-Tormo, M., Teichmann, S.A., and Vento-Tormo, R. (2020). CellPhoneDB: inferring cell-cell communication from combined expression of multi-subunit ligand-receptor complexes. Nat. Protoc. 15, 1484–1506.

38. Liu, D., Black, B.L., and Derynck, R. (2001). TGF-beta inhibits muscle differentiation through functional repression of myogenic transcription factors by Smad3. Genes Dev. 15, 2950–2966.

39. Brennan, T.J., Edmondson, D.G., Li, L., and Olson, E.N. (1991). Transforming growth factor beta represses the actions of myogenin through a mechanism independent of DNA binding. Proc. Natl. Acad. Sci. U. S. A. 88, 3822–3826.

40. De Micheli, A.J., Spector, J.A., Elemento, O., and Cosgrove, B.D. (2020). A reference single-cell transcriptomic atlas of human skeletal muscle tissue reveals bifurcated muscle stem cell populations. Skelet. Muscle 10, 19.

41. Jin, S., Guerrero-Juarez, C.F., Zhang, L., Chang, I., Ramos, R., Kuan, C.-H., Myung, P., Plikus, M.V., and Nie, Q. (2021). Inference and analysis of cell-cell communication using CellChat. Nat. Commun. 12, 1088.

42. Paliwal, P., Pishesha, N., Wijaya, D., and Conboy, I.M. (2012). Age dependent increase in the levels of osteopontin inhibits skeletal muscle regeneration. Aging 4, 553–566.

43. Juhas, M., Abutaleb, N., Wang, J.T., Ye, J., Shaikh, Z., Sriworarat, C., Qian, Y., and Bursac, N. (2018). Incorporation of macrophages into engineered skeletal muscle enables enhanced muscle regeneration. Nat Biomed Eng 2, 942–954.

44. Dokun, A.O., Keum, S., Hazarika, S., Li, Y., Lamonte, G.M., Wheeler, F., Marchuk, D.A., and Annex, B.H. (2008). A quantitative trait locus (LSq-1) on mouse chromosome 7 is linked to the absence of tissue loss after surgical hindlimb ischemia. Circulation 117, 1207–1215.

45. Picelli, S., Faridani, O.R., Björklund, A.K., Winberg, G., Sagasser, S., and Sandberg, R. (2014). Full-length RNA-seq from single cells using Smart-seq2. Nat. Protoc. 9, 171–181.

46. Liu, L., Cheung, T.H., Charville, G.W., and Rando, T.A. (2015). Isolation of skeletal muscle stem cells by fluorescence-activated cell sorting. Nat. Protoc. 10, 1612–1624.

47. Hao, Y., Hao, S., Andersen-Nissen, E., Mauck, W.M., 3rd, Zheng, S., Butler, A., Lee, M.J., Wilk, A.J., Darby, C., Zager, M., et al. (2021). Integrated analysis of multimodal single-cell data. Cell 184, 3573–3587.e29.

48. Korsunsky, I., Millard, N., Fan, J., Slowikowski, K., Zhang, F., Wei, K., Baglaenko, Y., Brenner, M., Loh, P.-R., and Raychaudhuri, S. (2019). Fast, sensitive and accurate integration of single-cell data with Harmony. Nat. Methods 16, 1289–1296.

49. Subramanian, A., Tamayo, P., Mootha, V.K., Mukherjee, S., Ebert, B.L., Gillette, M.A., Paulovich, A., Pomeroy, S.L., Golub, T.R., Lander, E.S., et al. (2005). Gene set enrichment analysis: a knowledge-based approach for interpreting genome-wide expression profiles. Proc. Natl. Acad. Sci. U. S. A. 102, 15545–15550.

50. Coates, P.J., Rundle, J.K., Lorimore, S.A., and Wright, E.G. (2008). Indirect macrophage responses to ionizing radiation: implications for genotype-dependent bystander signaling. Cancer Res. 68, 450–456.

51. Liberzon, A., Subramanian, A., Pinchback, R., Thorvaldsdóttir, H., Tamayo, P., and Mesirov, J.P. (2011). Molecular signatures database (MSigDB) 3.0. Bioinformatics 27, 1739–1740.

52. Cao, J., Spielmann, M., Qiu, X., Huang, X., Ibrahim, D.M., Hill, A.J., Zhang, F., Mundlos, S., Christiansen, L., Steemers, F.J., et al. (2019). The single-cell transcriptional landscape of mammalian organogenesis. Nature 566, 496–502.

53. Garcia-Alonso, L., Handfield, L.-F., Roberts, K., Nikolakopoulou, K., Fernando, R.C., Gardner, L., Woodhams, B., Arutyunyan, A., Polanski, K., Hoo, R., et al. (2021). Mapping the temporal and spatial dynamics of the human endometrium in vivo and in vitro. Nat. Genet. 53, 1698–1711.

54. Dobin, A., Davis, C.A., Schlesinger, F., Drenkow, J., Zaleski, C., Jha, S., Batut, P., Chaisson, M., and Gingeras, T.R. (2013). STAR: ultrafast universal RNA-seq aligner. Bioinformatics 29, 15–21.

55. Liao, Y., Smyth, G.K., and Shi, W. (2014). featureCounts: an efficient general purpose program for assigning sequence reads to genomic features. Bioinformatics 30, 923–930.

56. Sherman, B.T., Hao, M., Qiu, J., Jiao, X., Baseler, M.W., Lane, H.C., Imamichi, T., and Chang, W. (2022). DAVID: a web server for functional enrichment analysis and functional annotation of gene lists (2021 update). Nucleic Acids Res. 50, W216–W221.

57. Huang, D.W., Sherman, B.T., and Lempicki, R.A. (2009). Systematic and integrative analysis of large gene lists using DAVID bioinformatics resources. Nat. Protoc. 4, 44–57.

58. Ramírez, F., Ryan, D.P., Grüning, B., Bhardwaj, V., Kilpert, F., Richter, A.S., Heyne, S., Dündar, F., and Manke, T. (2016). deepTools2: a next generation web server for deep-sequencing data analysis. Nucleic Acids Res. 44, W160–W165.

59. Robinson, J.T., Thorvaldsdóttir, H., Winckler, W., Guttman, M., Lander, E.S., Getz, G., and Mesirov, J.P. (2011). Integrative genomics viewer. Nat. Biotechnol. 29, 24–26.

