## Supplementary figures and images for "Pro-inflammatory macrophages impair skeletal muscle regeneration in ischemic-damaged limbs by inducing precocious differentiation of satellite cells"

### Supplementary Figure 1

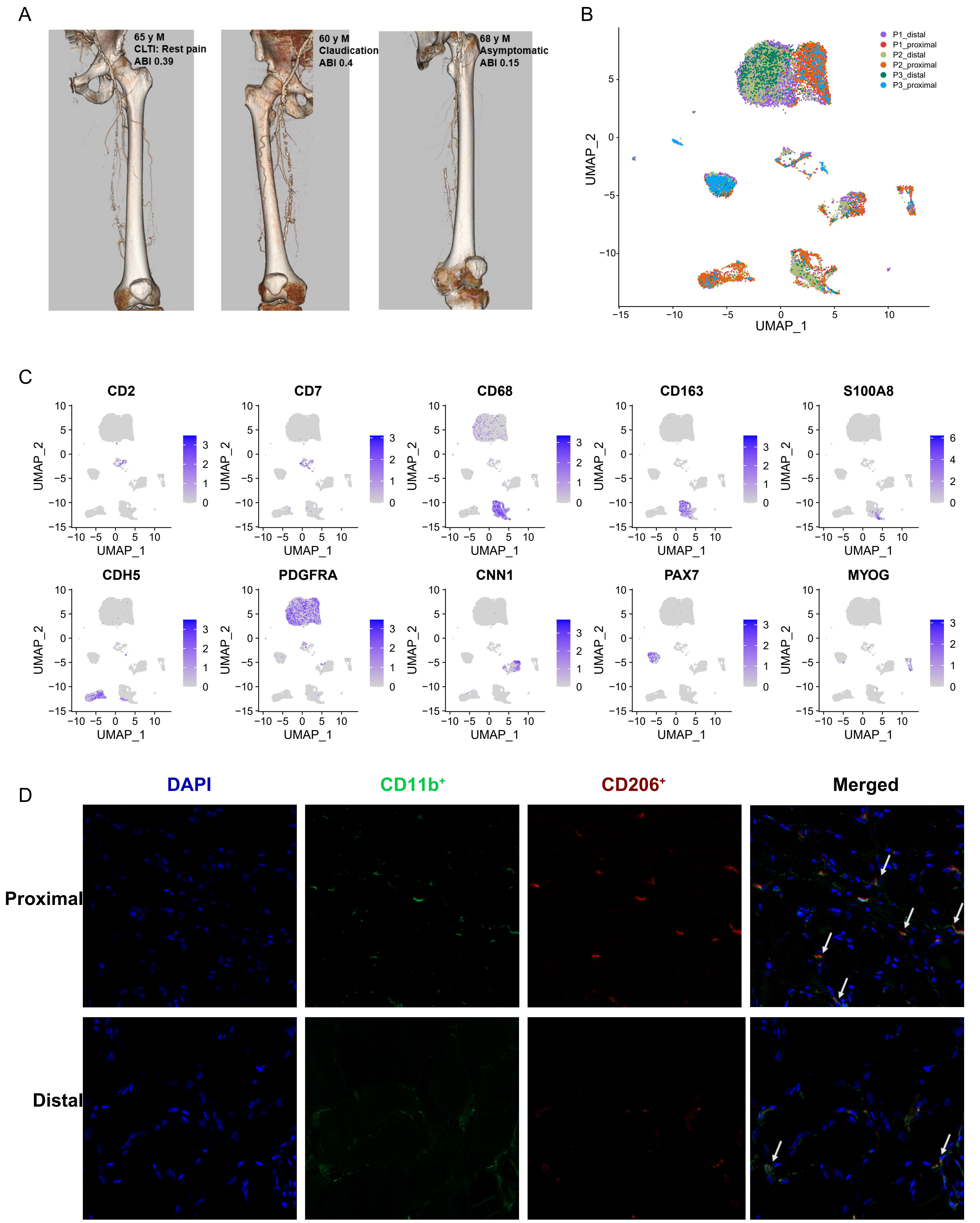

### Supplementary Figure 2

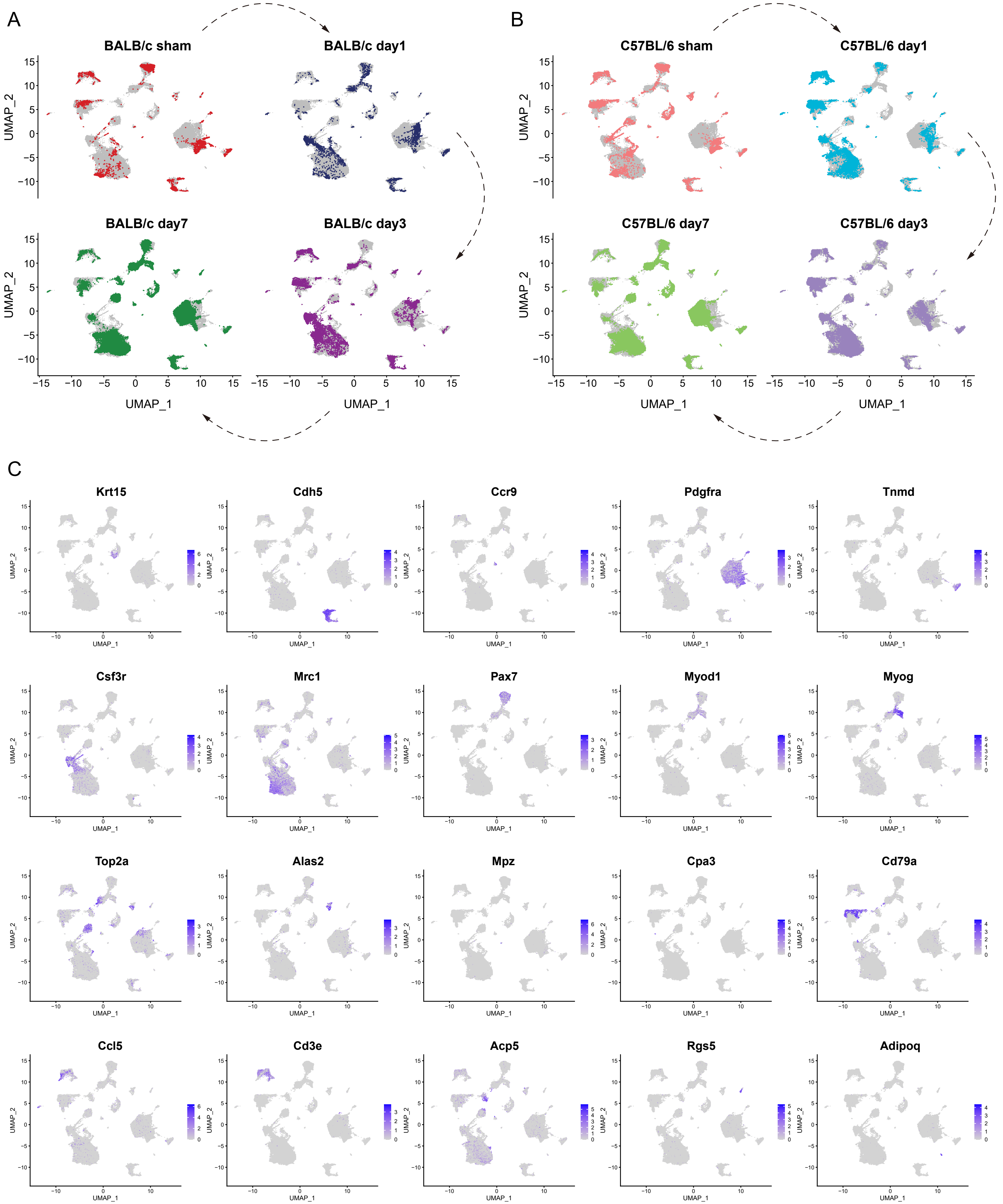

### Supplementary Figure 3

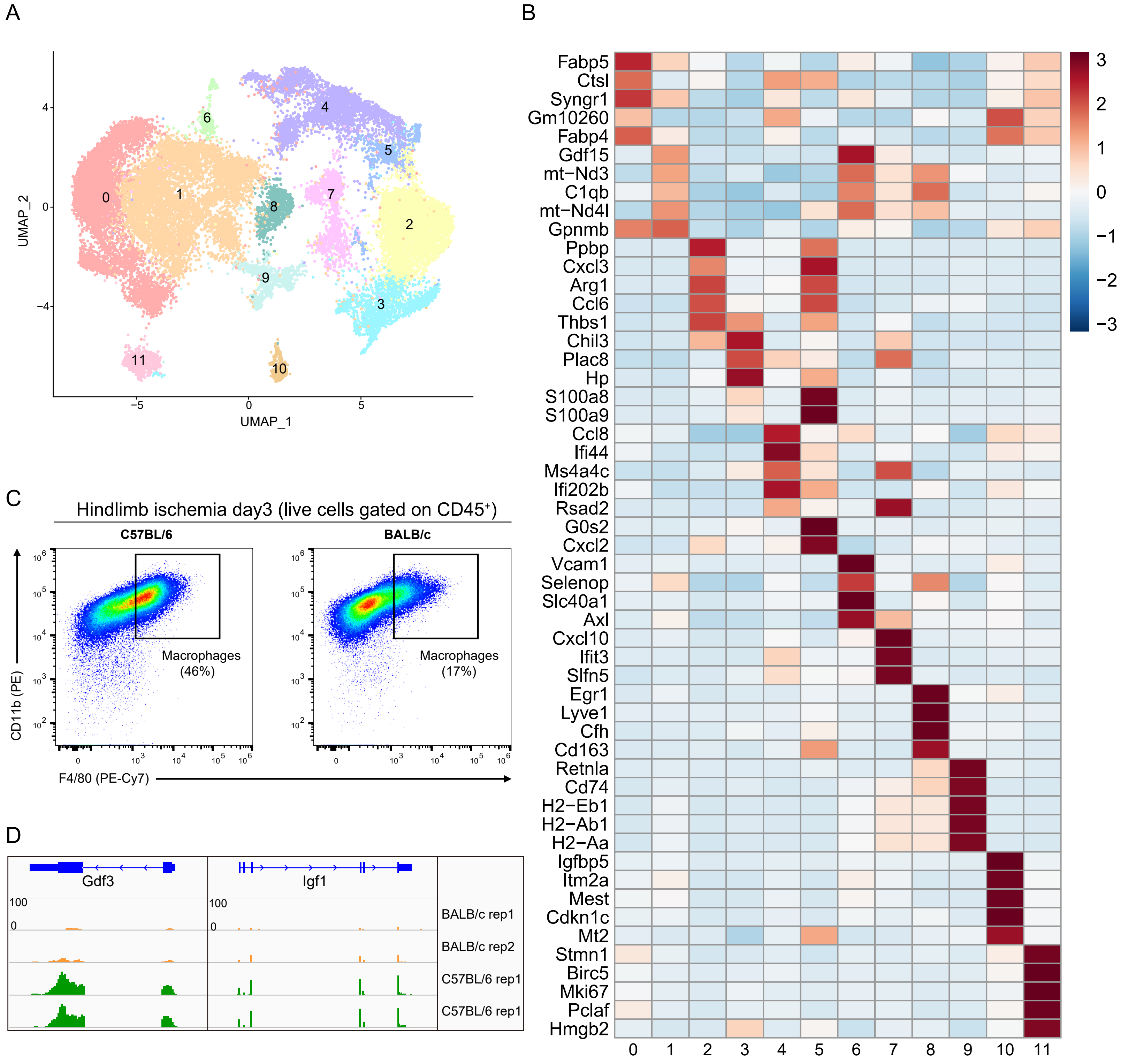

### Supplementary Figure 4

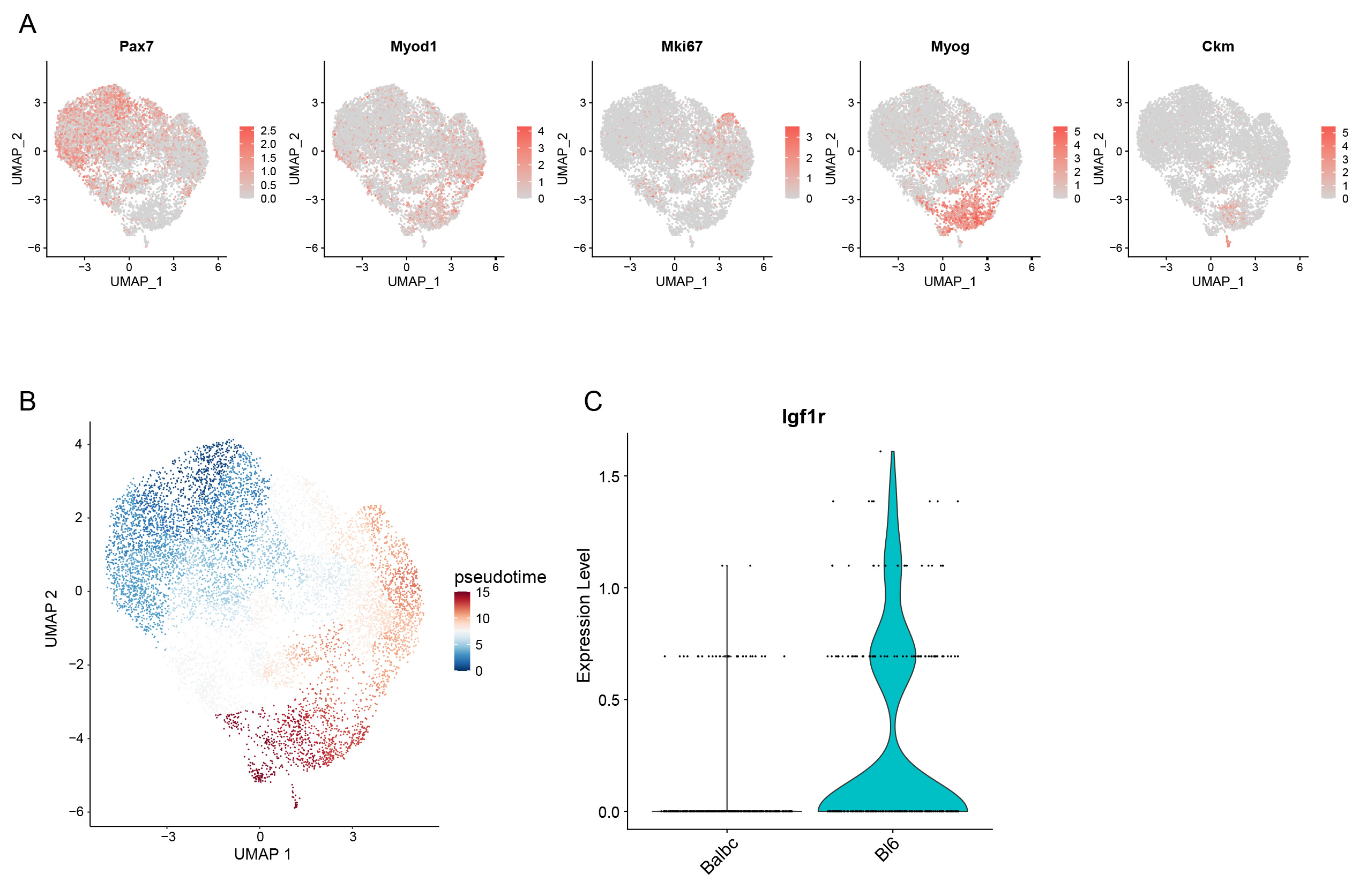

### Supplementary Figure 5

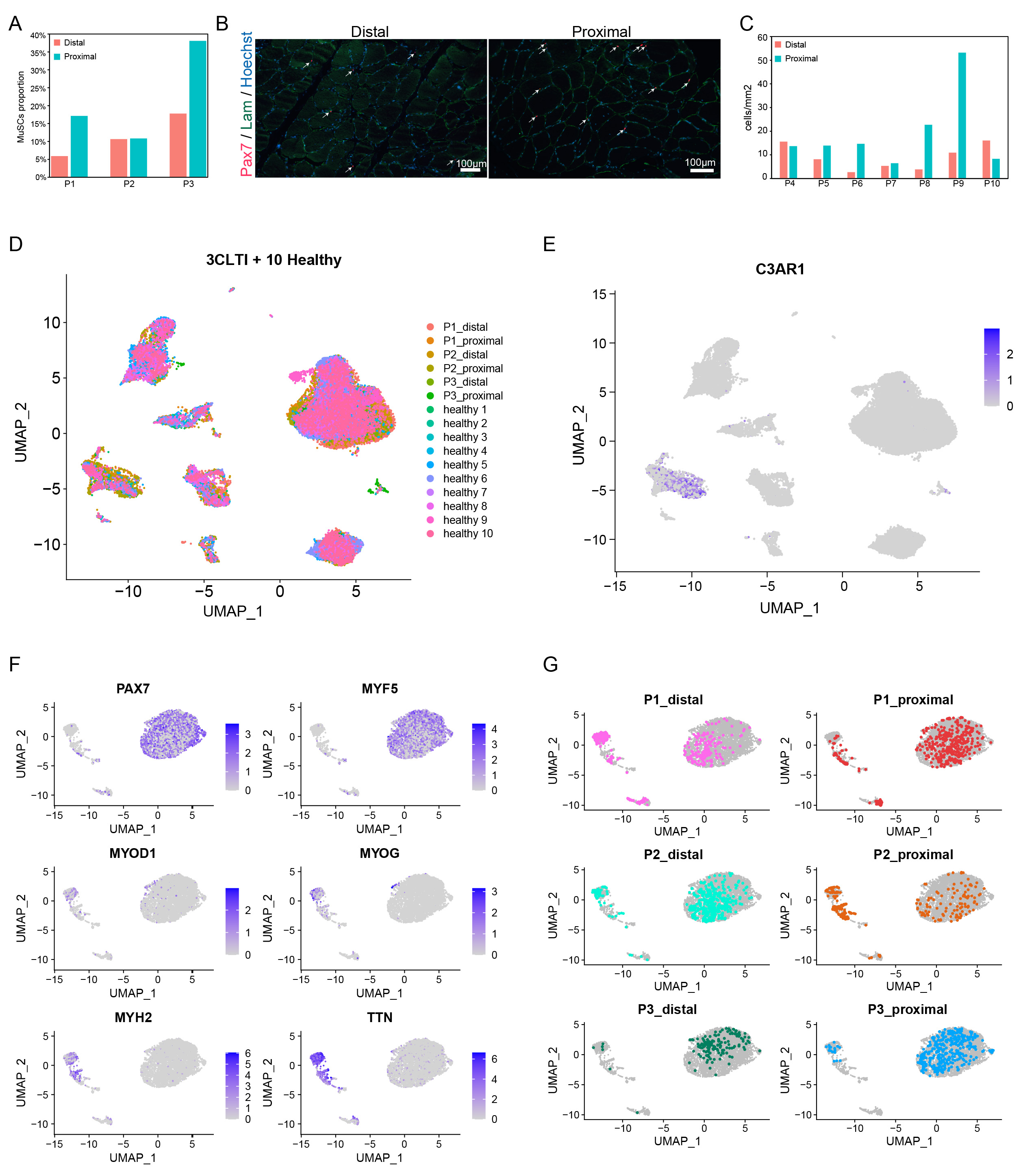
